# Label-free lymphocytes reconstitution using side scatter for optimal T cell manufacturing

**DOI:** 10.1101/2020.11.09.375584

**Authors:** Tongjin Wu, Yen Hoon Luah, Yongqiang Luo, Howard John Womersley, Lih Feng Cheow

## Abstract

Lymphocyte biology research commonly involves purification of lymphocyte subpopulations by fluorescence-activated cell sorting (FACS) or immunomagnetic separation (IMS), both of which typically rely on antibody labeling of validated cell markers. Methods enabling label-free segregation of lymphocyte subpopulations would be invaluable with regard to less-perturbation, simplicity and cost-effectiveness. Here, we introduce TRuST, a label-free approach for <u>T</u> cell <u>r</u>econstitution <u>u</u>sing <u>s</u>ide-scat<u>t</u>er (SSC). TRuST-sorted SSC^low^ cells enrich for CD4^+^ T and naïve T cells, while SSC^high^ cells enrich for CD8^+^ T, NK and differentiated T cells. Enrichment purity can be improved by computational gate design. SSC^low^ cells have superior expansion capacity and generate more central memory precursors with naïve-resembling cytokine responses. Moreover, we find that both T cell differentiation status and CD4/CD8 T ratio in the starting cellular material are critical attributes predicting T cell product quality and quantity. TRuST presents an effective and reliable technique for label-free lymphocytes selection and reconstitution.

## INTRODUCTION

Lymphocytes, mainly including T cells, natural killer (NK) cells and B cells, play an indispensable part in human immune system. Use of purified lymphocyte subsets for lymphocyte biology research is critical to dissect the functional heterogeneity of lymphocyte subpopulations. Fluorescence-activated cell sorting (FACS) (Cossarizza et al., 2019) and immunomagnetic separation (IMS) (Plouffe et al., 2015) are methods of choice to obtain purified cell types of interest, relying on the labeling with specific antibody conjugates such as fluorochrome-modified antibodies and magnetic beads-coupled antibodies. Despite powerful of these technologies, there still exist some limitations.

One of the most common concerns is antibody labeling-caused functional change of immune cells which are relatively sensitive to antibody-mediated activation/inhibition (Attanasio and Wherry, 2016; Chen and Flies, 2013), interfering with downstream cell biology studies. Thus, careful selection of validated antibody clones and strict experimental controls are generally required in experimental design. In addition, it is extremely difficult to release the bound fluorescent antibodies on cell surface if cell viability was to be maintained. These sorted cells are not suitable for re-labeling with the same fluorochrome conjugates due to occupancy of fluorescence channels, resulting in less flexibility in the choice of fluorescent antibodies for multiparametric flow cytometry experiments. Also, these cells might not be feasible for downstream cell culture assays if the occupied target markers were necessary to be re-engaged for cell functionality. For example, the labeling of CD3 receptor for T cells enrichment by FACS might affect the activation/expansion of T cells when CD3/CD28 engagement is required for cell stimulation (Roddie et al., 2019). Superparamagnetic particles-antibody coupling technique combined with magnetic isolation is shown to be capable of minimizing the effects introduced by fluorescent antibodies occupancy thanks to its effective separation with only unsaturated concentration of bead-antibody conjugates (Grutzkau and Radbruch, 2010; Plouffe et al., 2015). However, concerns arising from the use of nanosized magnetic beads include internalization of magnetic beads and preservation of magnetic properties for an extended period in cells that are subjected to limited proliferation, which calls for caution when repetitive positive isolation steps after in vitro culture are required (Laghmouchi et al., 2020; Teeman et al., 2019). More recently, Miltenyi Biotec has introduced the REAlease Fluorochrome Technology that relies on engineered antibody fragments with low epitope binding affinities which allows stained antibodies to be released. Design of releasable superparamagnetic beads-antibody conjugate could be of interest to improve magnetism-based cell separation. Nevertheless, the time and cost for sample processing are inevitably increased and concerns relating to cell labeling-caused alteration on downstream assays still exist. Therefore, with regard to less perturbation on target cells and enhanced flexibility in sample processing, approaches that enable label-free cell type separation from mixed lymphocytes would be highly attractive and great useful in lymphocyte biology studies.

Selection of specific T cell types has also seen its utility in the field of cellular immunotherapy. Insight into attributes of therapeutic potency based on T cell receptor (TCR)-redirected T cells such as chimeric antigen receptor (CAR) T cells and TCR-T cells is a central goal of T cell immunotherapy (Majzner and Mackall, 2019; Rafiq et al., 2020). The quality of T cell products generated ex vivo has been recognized as a critical factor affecting immunotherapy efficacy (Deng et al., 2020; Finney et al., 2019; Fraietta et al., 2018; Roddie et al., 2019). Unfractionated peripheral blood mononuclear cell (PBMC) concentrates are commonly used for T cell production, leading to uncontrollable batch-to-batch variation. Separate expansion of CD4^+^ T and CD8^+^ T cells pre-enriched by immunomagnetic selection enhances product consistency and manufacturing feasibility (Gardner et al., 2017; Shah et al., 2020; Sommermeyer et al., 2016; Turtle et al., 2016). Of note, CD4^+^ T cells help is critical for shaping the adaptive immunity of CD8^+^ T cells in vivo (Nakanishi et al., 2009; Sun et al., 2004) and improves CD8^+^ T cells tumor-eradicating potency (Alspach et al., 2019; Arina et al., 2017; Nakanishi et al., 2009; Schietinger et al., 2010). However, the notion, whether CD4^+^ T and CD8^+^ T cells should be expanded separately or together at designed ratio in vitro, especially in the scenario where external antigens are absent, remains to be clarified. In addition, adoptive transfer of T cell products derived from less-differentiated T cell subsets shows better curative potential than their more-differentiated counterparts as observed in both animal works (Berger et al., 2008; Gattinoni et al., 2011; Hinrichs et al., 2009; Hinrichs et al., 2011; Wang et al., 2011) and clinical studies (Finney et al., 2019; Fraietta et al., 2018; Xu et al., 2014). These studies suggest that preselecting specific T cell subtypes would be very useful for cell manufacturing aiming to improve immunotherapy efficacy. Undoubtedly, increased demand for therapeutic T cell product calls for cost-effectiveness and manufacturing simplicity. Methods enabling the segregation of ideal cell subsets without the use of costly antibodies would be invaluable by omitting the requirement of large amount of clinical grade immunomagnetic antibodies and complicated downstream processing workflows.

There has been an increasing number of thought-provoking studies on label-free lymphocytes identification. Light scattering, a parameter commonly used in flow cytometry, is capable of measuring cell size via forward scatter (FSC) and cellular granularity by side scatter (SSC) (Cossarizza et al., 2019). Light scattering has already seen its versatile applications in the screening of lymphocyte samples to distinguish healthy and disease patients such us B-cell chronic lymphocytic leukemia (B-CLL) (Terstappen et al., 1988), cutaneous T-cell lymphoma (Clark et al., 2011) and chronic hepatitises B or C infection (Ruban et al., 2010). The principle underlining light scattering relied disease diagnostics could be that, in general, CD8^+^ T cells has higher SSC than CD4^+^ T cells while B cells tend to locate between the resting and activated CD8^+^ T cells populations (Terstappen et al., 1986a; Terstappen et al., 1986b). Of note, most of the reports have not clearly clarified the potential confounding effects resulting from cell size on SSC pattern given that unhealthy or activated lymphocytes with altered size (higher FSC) are likely to change their SSC profile also (Clark et al., 2011; Loudon et al., 1988; Terstappen et al., 1988). Thus, the question arising is whether SSC distribution is enough to tell the differences of lymphocyte cell types that have indistinguishable cell size, for example, healthy lymphocyte subpopulations under resting status. Moreover, to date, a closer look at the subtle scattering distribution of T cell differentiation lineages (e.g. naïve/memory T cells) has been neglected. Memory/effector T cells are observed to have more granules than their naïve counterparts under transmission electron microscope (Dimeloe et al., 2016). These findings imply that cell general biophysical attributes (e.g. light scattering) may be further exploited to select for lymphocyte subpopulations with particular functionality, such as for immunotherapy.

Taken together, these inadequacies highlight the need for new methods capable of selecting lymphocyte subpopulations in a label-free manner by relying on general cell biophysical properties. To achieve this, we initially had an in-depth analysis of the light scattering profile (FSC/SSC) of resting human lymphocyte subsets, including NK cells, B cells, total CD4^+^ T cells, total CD8^+^ T cells, as well as CD4^+^ or CD8^+^ naïve/memory T cell subsets. We found that SSC instead of FSC is a robust biophysical property to distinguish lymphocyte subsets. Generally, lymphocytes with “helper” function such as CD4^+^ T cells and B cells have a lower SSC than that with “cytotoxic” function such as CD8^+^ T cells and NK cells. Interestingly, this discrepancy in SSC profile is even more apparent among T cell differentiation lineages regardless of CD4^+^ or CD8^+^ T cell type. Younger T cells such as naïve T cells and memory precursors have a significantly lower SSC than their more-differentiated counterparts including central memory T cells, effector memory T cells and terminally effector memory T cells. Inspired by these findings, we developed TRuST, a label-free method for T cell reconstitution using side scatter. TRuST-relied T cell segregation is based on two gating strategies. One is computational gate search aimed to maximize the purity of target cell type of interest. The other is manual gate selection which is easier and more flexible in reconstituting lymphocyte composition instead of focusing on single cell subset. We further demonstrated that reconstituted SSC^low^ cells that are enriched for CD4^+^ T and naïve T cells have superior expansion capacity and produce more central memory precursors. Finally, using the TRuST-based T cell segregation, we were able to find that both less-differentiated T cell subpopulations and high CD4^+^ T to CD8^+^ T ratio in starting cellular material are critical attributes predicting T cell product quality and quantity. This study established TRuST as an effective and reliable technique for label-free lymphocytes selection and reconstitution, which would be useful for lymphocytes biology studies and T cell manufacturing for improved immunotherapy.

## RESULTS

### TRuST development

TRuST is a label-free approach for lymphocyte subpopulations selection and T cell reconstitution. The development of TRuST was inspired by the intrinsic side-scattering difference of lymphocyte subpopulations including CD4^+^ T cells, CD8^+^ T cells, NK cells, B cells, and main T cell differentiation lineages including naïve (T_N_, CD45RA^high^ CCR7^high^), central memory (T_CM_, CD45RA^dim/−^ CCR7^low^), effector memory (T_EM_, CD45RA^dim/−^ CCR7^−^) and terminally differentiated effector (T_EMRA_, CD45RA^high^ CCR7^−^) (**Figures 1A, S1A, and S1B**) (Bohler et al., 2007; Seder and Ahmed, 2003). Due to the heterogeneous SSC patterns of lymphocyte subpopulations, using the pre-stained lymphocytes for each new sample, it is feasible to find the optimum gate that has the desired purity and yield for target cells of interest via GateID, a computational gate search algorithm for cell type purification (Baron et al., 2019) (**Figure 1B**). Alternatively, a series of manual gates can be set to determine the optimum gate for T cell reconstitution depending on the research aims (**Figure 1C**). The selected gate can be simply adopted and applied for the sorting of unstained lymphocytes (**Figure 1D**). For downstream cell culture assays such as T cell expansion for immunotherapy, the total lymphocytes can be directly sorted into SSC^low^ and SSC^high^ groups that consist of functionally distinct T cell subsets. The sorted cells are subjected to canonical expansion protocol using anti-CD3/CD28 microbeads plus interleukin-2 (IL-2) (**Figures 1D and 1E**). This enables us to study how the initial cell composition affect T cell expansion dynamics, phenotypes formation, and functionality shaping in multifunctional cytokine expression (**Figures 1E and S1C**), and the attributes that contribute to the discrepancies of final T cell products (**Figure 1E**). Together, this work flow demonstrates the utility of TRuST for label-free lymphocyte selection and T cell constitution for downstream lymphocytes biology study.

**Figure 1.**
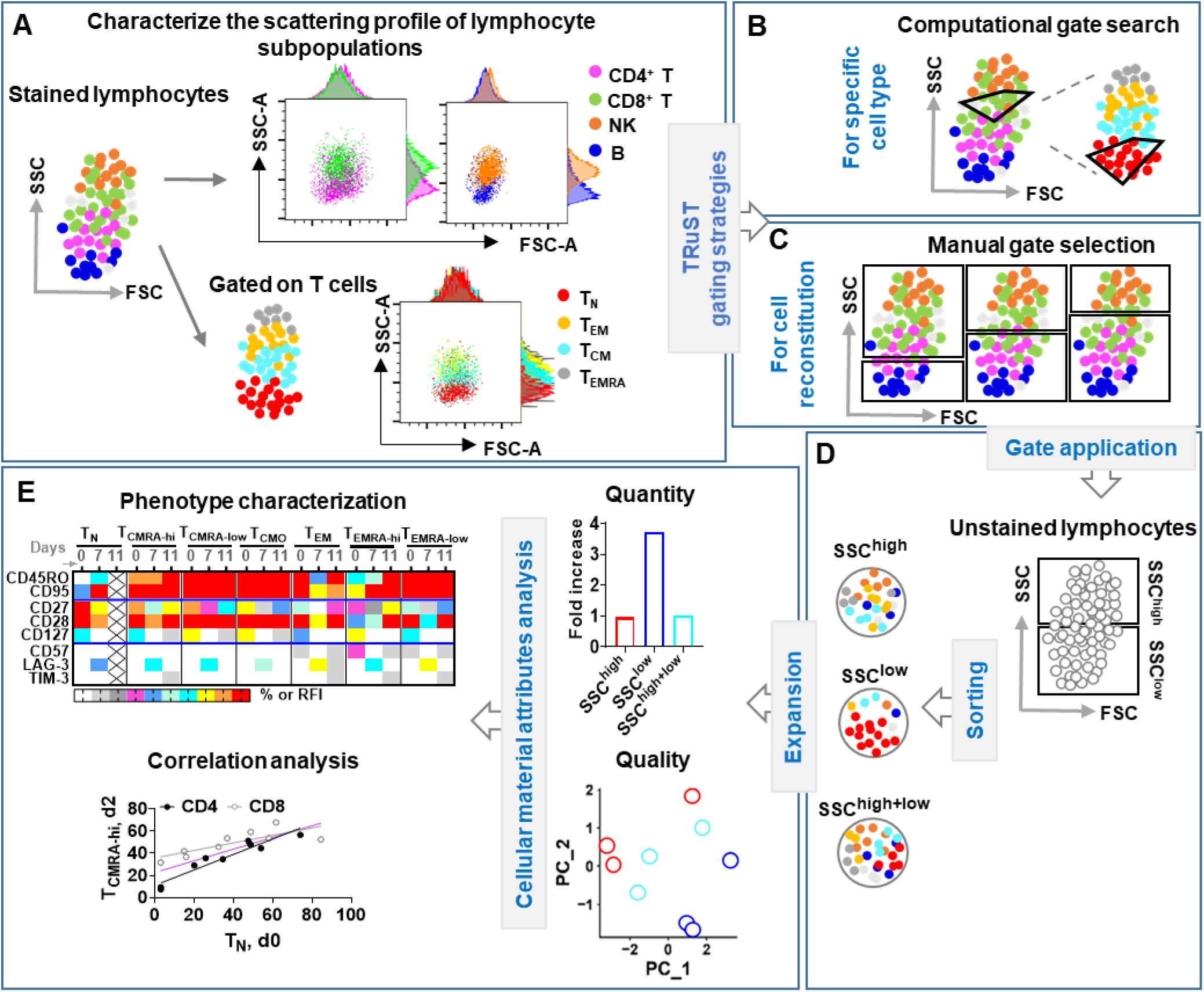
TRuST: An effective approach for label-free T cell selection and reconstitution. (**A**) PBMC samples were stained with lymphocyte subpopulation-representing fluorescence antibodies. Data was acquired by flow cytometer and the forward/side-scattering (FSC/SSC) distribution of main lymphocyte populations and T cell naïve/memory subsets were studied. (**B**) Based on the FSC/SSC profile of lymphocyte populations, a computational gate design algorithm was performed to enrich specific target cell type with optimal purity and yield. (**C**) Alternatively, a manual gating strategy relied on the SSC intensity was used to reconstitute lymphocyte populations with distinct functionality. (**D**) The manually selected gate was applied for the sorting of unstained lymphocytes for optimum T cell production. (**E**) A comprehensive comparison of the quantity and quality of T cell products that were derived from cellular materials with different SSC intensity. Characterization of factors contributing to the distinct products by longitudinally phenotypic assessment and correlation analysis. See also **Table S1** and **Methods**.

### TRuST enables effective reconstitution of functionally distinct lymphocyte subpopulations

Lymphocyte cluster can be robustly separated from monocytes and granulocytes based on their FSC/SSC distribution as has been routinely used in flow cytometry. However, a comprehensive study of the subtle scattering profile within various lymphocyte subpopulations is still absent (**Figure 1A**). Using human lymphocytes pre-stained with cell type markers, we found that CD4^+^ T, B cells and T_N_ have a lower SSC (low granularity) compared to CD8^+^ T, NK and differentiated T cells such as T_CM_, T_EM_ and T_EMRA_ (high granularity) (**Figures 2A and 2B**). With the gradual increase of SSC intensity, there are more CD8^+^ T and NK cells, as well as more differentiated T cell subsets, leading to percentage reduction of CD4^+^ T and T_N_ cells (**Figure S2**). For consistency throughout this work, we manually dived the total lymphocytes into two equal parts (SSC^high^ and SSC^low^). This gating method is robust enough to either sample staining procedures (**Figures S3A and S3B**) or voltages change applied during flow cytometry data acquiring (**Figures S3A, S3C, and S3D**). On the contrary, FSC can only differentiate B cells from NK cells (**Figures S3E-S3G**). The nearly unaltered light scattering profiles between stained and unstained cells (**Figure S3B**) enable us to directly apply the gate in sorting of unstained lymphocytes (**Figure 2C**). Similarly, sorted SSC^low^ group enriches for CD4^+^ T cells and naïve T cells while SSC^high^ cluster enriches for CD8^+^ T, NK and other more-differentiated T cell counterparts including T_CM_, T_EM_ and T_EMRA_ (**Figures 2D–2G**). Deep-ultraviolet light (UV) has been adopted for label-free molecular imaging (Zeskind et al., 2007) and hematology analysis (Ojaghi et al., 2020) thanks to its shorter wavelength. Thus, it is interesting to study whether UV light (e.g. UV405 nm) can be used to further improve the scattering resolution of cell morphometric structure under blue light (e.g. B488 nm). However, we observed that a double side-scatter (UV405/B488) based sorting strategy (**Figure S3H**) did not improve the general segregation efficiency of CD4/CD8 T cells, NK/B cells and naïve/memory T cells as well (**Figures S3I-S3M**). Therefore, global cellular granularity, as represented by SSC intensity, could be a general biophysical property used for the separation of lymphocyte subpopulations.

**Figure 2.**
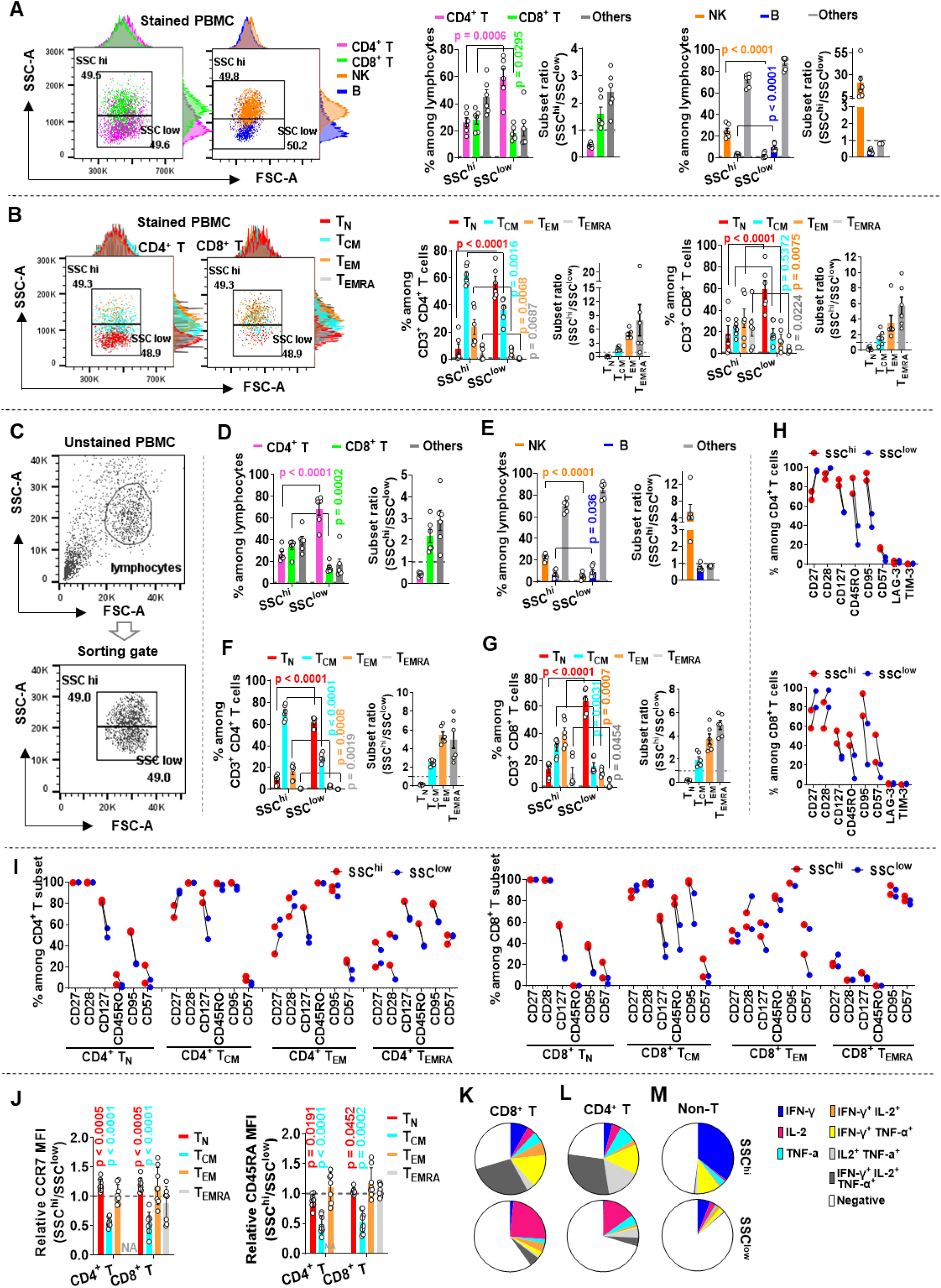
SSC is a robust biophysical property for TRuST development. (**A and B**) PBMC samples were stained with surface marker antibodies before flow cytometry data acquiring. Dot plots are representative FSC/SSC profiles of CD3^+^ CD4^+^ T/CD3^+^ CD8^+^ T cells (**A, left dot plot**), NK/B cells (**A, right dot plot**), and T_N_/T_CM_/T_EM_/T_EMRA_ cell subsets (**B**). For statistical analysis, samples were gated to equally divide the cells into two clusters based on the cluster of total lymphocytes (not shown in the dot plots) (**A and B, dot plots**) and the percentage of CD3^+^ CD4^+^ T/CD3^+^ CD8^+^ T cells, NK/B cells, and T_N_/T_CM_/T_EM_/T_EMRA_ cell subsets within SSC^hi^ or SSC^low^ cluster were summarized (**A and B, histograms**). Two independent experiments for each donor (n = 3) were compiled. A two-tailed, paired *t*-test between the two groups was performed. Error bars indicate mean ± SEM. (**C-G**) Unstained PBMC samples were equally sorted into SSC^hi^ and SSC^low^ populations before lymphocyte subtypes determination (**C**). Characterization of the percentage of CD3^+^ CD4^+^ T/CD3^+^ CD8^+^ T cells **(D**), NK/B cells (**E**), and T_N_/T_CM_/T_EM_/T_EMRA_ (**F and G**) among the sorted SSC^hi^ and SSC^low^ cell populations. Data from three donor samples with two independent experiments for each donor were compiled. A two-tailed, paired *t*-test between the two groups was performed. Error bars indicate mean ± SEM. **(H and I)** A comparison of the expression profiles of selected phenotyping markers in sorted SSC^hi^ and SSC^low^ cells (**H**), and characterization of the phenotypic traits of the same naïve/memory T subset derived from SSC^hi^ or SSC^low^ cell clusters (**I**). Each dot represents data of one donor PBMC (**H and I**). **(J)** The relative expression level of CCR7 and CD45RA in CD4^+^ T or CD8^+^ T naïve/memory subsets from sorted SSC^hi^ and SSC^low^ cells. Plots represent data from three donor PBMC. Cell subset nearly undetectable is indicated as “NA”. A two-tailed, unpaired *t*-test between SSC^hi^ and SSC^low^ groups was performed. Error bars indicate mean ± SEM. (**K-M**) Sorted SSC^hi^ and SSC^low^ cells were stimulated with PMA plus ionomycin for 4 hours before intracellular staining. Shown are the multifunctional cytokine expression profile of CD3^+^ CD8^+^ T cells (**K**), CD3^+^ CD8-T cells (**L**), and CD3-cells (**M**) within sorted sorted SSC^hi^ or SSC^low^ cells (n = 2). See also **Figure S1** for gating methods in determining cell types and multifunctionality analysis, **Figures S2** and **S3** for manual gating selection.

Next, we sought to characterize the phenotypes of sorted cell populations based solely on SSC intensity. The sorted SSC^low^ cells displayed a higher expression of costimulatory receptors such as CD27 and CD28 but a lower expression of differentiation marker CD127, memory marker CD45RO and activation/inhibition factors CD95/CD57 (**Figure 2H**), consistent with a larger number of naïve T cells in this cell cluster. Unexpectedly, T cells of the same differentiation lineage but different SSC intensity also manifested some distinguishable phenotypic traits (**Figures 2I and 2J**). For example, either T_N_ or T_CM_ in SSC^low^ group had reduced CD127, CD95 and to a less extent CD57 (**Figure 2I**). Instead, T_CM_ of SSC^low^ cluster had increased CD27 (**Figure 2I**) and CCR7/CD45RA abundance (**Figure 2J**), suggesting its central memory precursor-like phenotype. Functionally, in response to short pulse of protein kinase C (PKC) stimulus, the SSC^low^ lymphocytes produced a large amount of IL-2 but reduced IFN-γ-involved immediate effector cytokines (**Figures 2K and 2L**), resembling cytokine expression dynamics of early-differentiated T cells (Denton et al., 2011; Hinrichs et al., 2009). The large amount of IFN-γ but near absence of IL-2 expression in non-T lymphocytes of SSC^high^ group is in line with a high proportion of NK cells in this cluster (**Figures 2M and 2E**). Taken together, these findings suggest that younger T cell lineages including naïve T cells and central memory precursors are intrinsically endowed with a lower SSC, which, concurrently with a higher ratio of CD4^+^ T to CD8^+^ T cells can be effectively enriched by TRuST.

### TRuST enhances lymphocyte subset selection using computational gate design

While manual gating for cell sorting is a simple and effective way to reconstitute lymphocytes, it could be less efficient for the selection of specific lymphocyte subset. Thus, we took advantage of the recently developed computational algorithm, GateID (Baron et al., 2019), that relies on the general properties (e.g. cell size, granularity and mitochondrial content) to search for the optimum gate for cell type enrichment. Using this method, it is possible to obtain lymphocyte subsets of interest with desired purity and yield (**Figure 3**).

**Figure 3.**
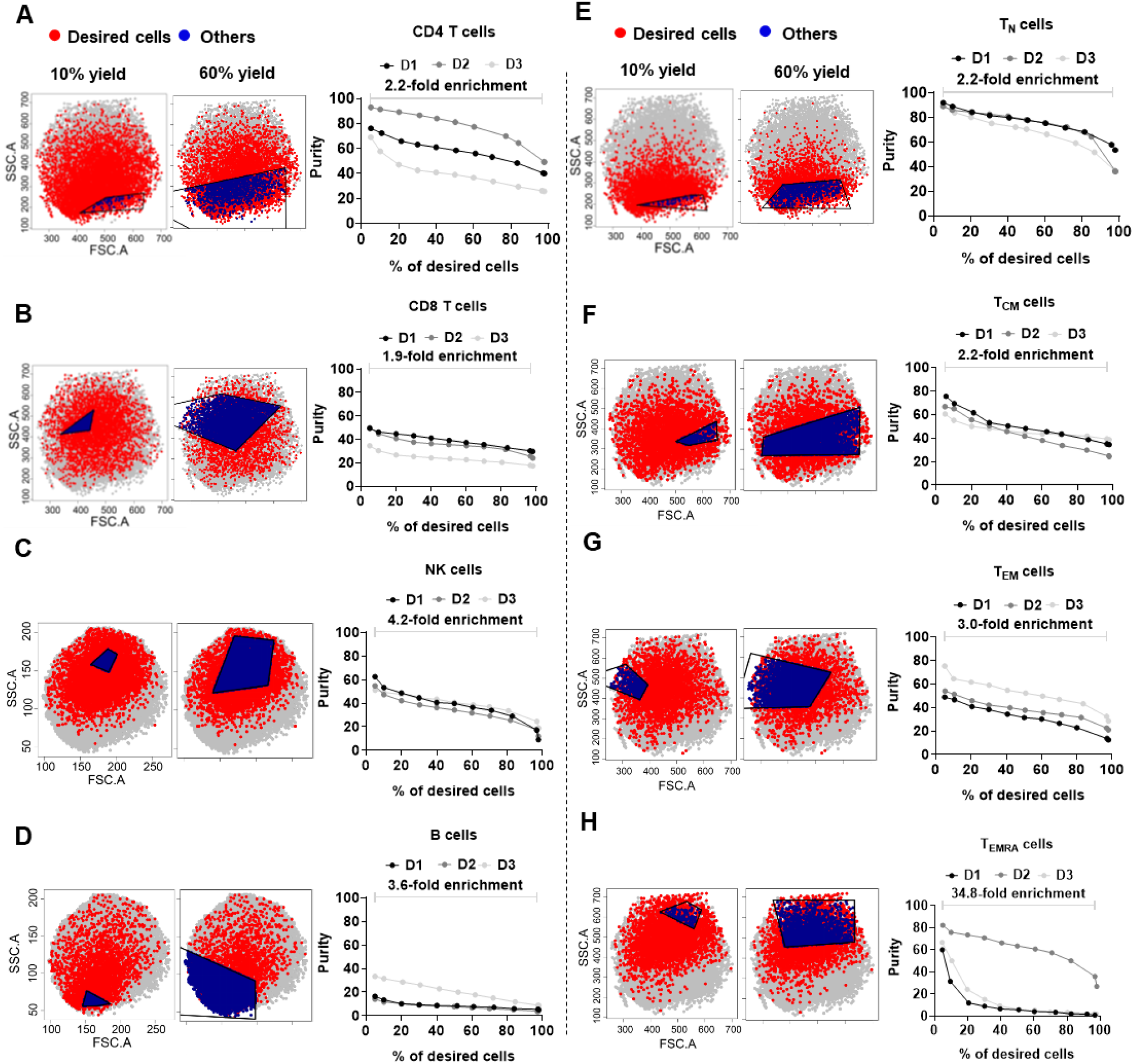
TRuST enhances enrichment of specific lymphocyte subset using computational gate design. (**A-H**) The raw readouts corresponding to FSC and SSC parameters for each cell type as shown in Figures 2A and 2B were exported, named and reanalyzed by GateID algorithm to select the desired cell type. The enrichment efficiency of CD3^+^ CD4^+^ T (**A**), CD3^+^ CD8^+^ T (**B**), NK (**C**) and B cells (**D**), or selection of total T_N_ (**E**), T_CM_ (**F**), T_EM_ (**G**) and T_EMRA_ subsets (**H**) were plotted by automatically adjustable gating threshold. For each cell type/subset, two representative gating plots corresponding to 10% and 60% yields of desired cells, and the correlation plots between purity and yield are shown. An average enrichment fold-increase between ~100% yield to ~5% yield (n = 3) is indicated. The percentage of desired cells (yield) is calculated as the number of desired cells within the gating to that of total desired cells among the whole cell population. The purity indicates the proportion of desired cells to that of total cell numbers within each gate. See also **Methods**.

Naïve T cells can be obtained with the highest purity and acceptable yield (**Figure 3E**), in line with their location within the cell group of lowest SSC (**Figure 2B**). T_CM_, T_EM_, T_EMRA_, total CD4^+^ T and NK cells can also be significantly enriched, but at the expense of quickly dropped yields (**Figures 3F–3H, 3A, and 3C**). On the other hand, CD8^+^ T cells and B cells tend to be the most difficult lymphocytes to be enriched with an improved purify and appreciable yield, partially due to their relatively homogeneous side-scattering distribution diffused into other lymphocytes (**Figures 3B and 3D**). Notably, given that a comparable FSC/SSC pattern was observed between unstained and stained cell samples (**Figure S3B**), it would be possible to directly apply the predicted gate in actual sorting experiments without gate normalization. Thus, combining TRuST with computational gate search, it is feasible to improve the purity of lymphocyte subsets with a reduced yet still acceptable recovery rate.

### TRuST-relied lymphocytes reconstitution improves T cell product quantity and quality

When practical feasibility and manufacturing cost are taken into consideration, such as T cell production for adoptive immunotherapy, it might not be necessary to enrich for a very specific cell type at the expense of greatly reduced cell recovery. Instead, strategies that enable the reconstitution of lymphocyte subpopulations to maximally improve cell production without compromising yield and quality would be preferable.

Given that T cell product derived from less-differentiated T cells show better outcome in adoptive immunotherapy (Berger et al., 2008; Finney et al., 2019; Fraietta et al., 2018; Hinrichs et al., 2011), we next investigated how compositional difference of starting T cell materials reconstituted solely by TRuST under manual gate selection (**Figure 2C**) may affect the diversity of final T cell products. In response to anti-CD3/CD28 microbeads stimulation, an antigen-presenting cells-mimic activation approach widely used for T cell manufacturing, SSC^low^ cells consisting of T_N_ and younger T_CM_ cells were endowed with superior proliferation capacity, and achieved an average increase of total nucleated cells (TNC) by 2.3-fold at day 7 and 3.2-fold at day 11 with 3-fold increase for CD4^+^ T and 4-fold increase for CD8^+^ T subtypes (**Figures 4A–4C and S4A-S4C**). Unexpectedly, the expansion potential of the SSC^high+low^ group appeared to be equivalent to that of SSC^high^ population (**Figures 4A–4C**), likely due to the precocious differentiation of naïve cells driven by existing effector/memory cells (Klebanoff et al., 2016) and the inhibitory effects caused by large amount of NK cells (Cook and Whitmire, 2013).

**Figure 4.**
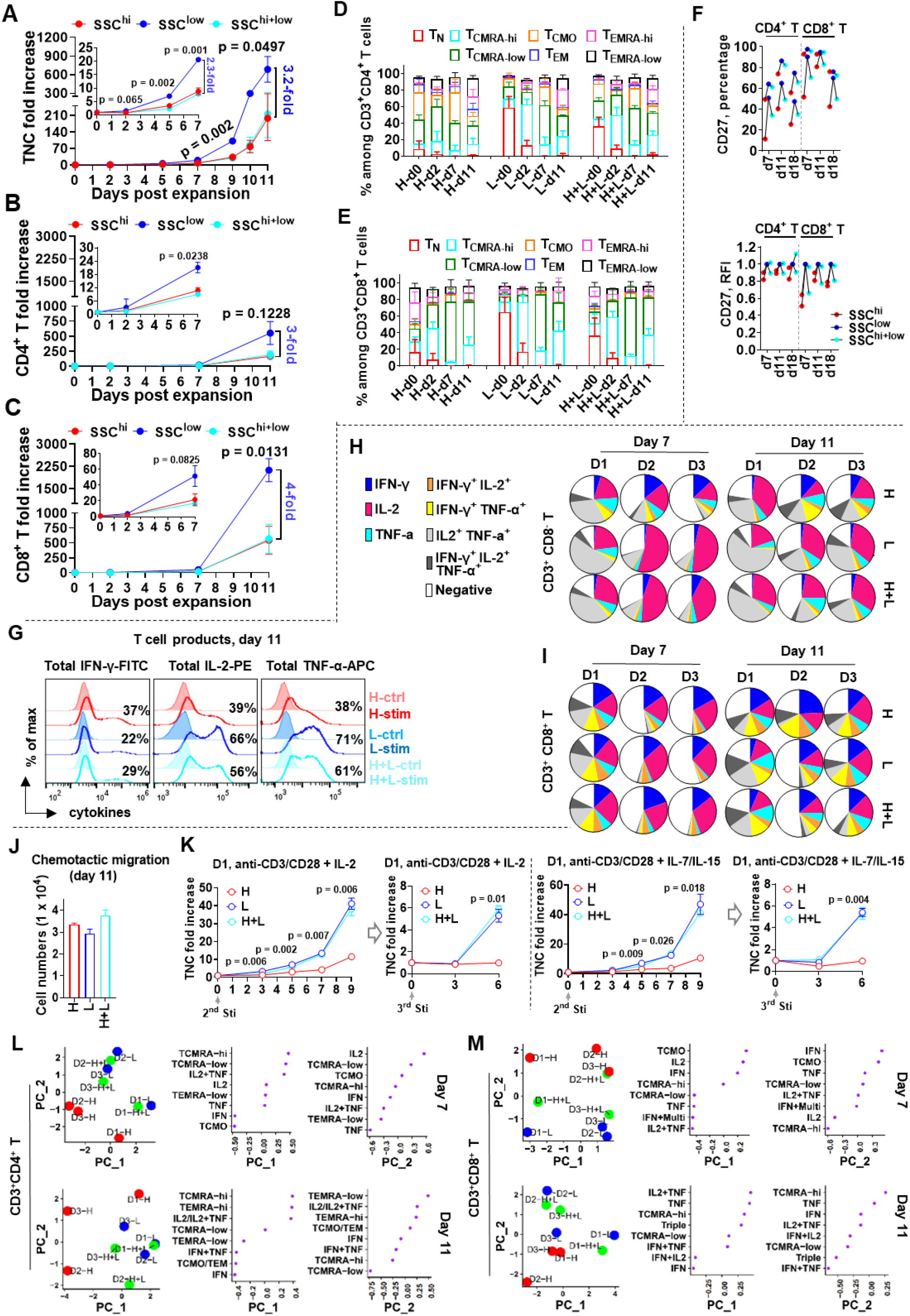
TRuST-relied lymphocytes reconstitution improves T cell product quantity and quality. (**A-C**) Expansion efficiency of lymphocytes derived from SSC^hi^, SSC^low^, or reconstituted SSC^hi^+low cells at a ratio of 1:1 are represented by TNC fold increase (**A**), CD3^+^ CD4^+^ T (**B**) and CD3^+^ CD8^+^ T fold increase (**C**). Data are presented as means ± S.E.M (n = 3). Paired two-tailed *t*-test between SSC^low^ and SSC^hi^ group was performed. Error bars indicate mean ± SEM. (**D and E**) Compositional change of naive/memory subsets in CD3^+^ CD4^+^ T (**D**) or CD3^+^ CD8^+^ T cells (**E**) plotted along the days of expansion. Error bars indicate mean ± SEM. (**F**) Percentage and relative fluorescence intensity (RFI) of CD27. (**G-I**) Representative histogram plot of IFN-γ/IL-2/TNF-α expression of cell product at day 11 post expansion derived from cell groups with different side-scattering intensity (**G**). Multifunctional cytokine expression of total CD4^+^ T (**H**) or CD8^+^ T (**I**) cells at day 7 and day 11 post expansion (n = 3) with 2 independent experiments for donor 1 and 2. Migration capacity of T cells at day 11 post expansion derived from SSC^hi^, SSC^low^ and SSC^hi^+low groups. Error bars indicate mean ± SEM. Responsiveness of effector cells descended from SSC^hi^, SSC^low^ and SSC^hi^+low groups at day 11 post expansion to multi-stimulation by CD3/CD28 engagement with IL-2 (**top**) or homeostatic cytokines IL-7 plus IL-15 (**bottom**). Duplicates for each sample were performed. Unpaired two-tailed *t*-test between SSC^low^ and SSC^hi^ group was performed. Error bars indicate mean ± SEM. (**L and M**) All phenotypic parameters and cytokine-expressing values were integrated together for PCA calculation. For data from day 7, IFN-γ^+^ IL-2^+^/ IFN-γ^+^ TNF-α^+^/ IFN-γ^+^ IL-2^+^ TNF-α^+^ were combined as “IFN+Multi”. “H” and “L” indicate “SSC^hi^” and “SSC^low^”, respectively. See also **Figures S4A-4C and S4E-4G** for individual data, and **Figure S4D** for gating method in cell differential lineages characterization.

Yellow fever-specific central memory CD8^+^ T cells with high CD45RA expression was demonstrated to be a critical cell subpopulation accounting for persistent immune memory, which resembled stem cell-like memory subset (CCR7^+^ CD45RA^+^ CD58^+^ CD95^+^ CD28^+^ CD27^+^s T_SCM_) (Fuertes Marraco et al., 2015). Thus, based on the dynamic change of CCR7/CD45RA expression, an in-depth gating strategy was used to classify T cell subsets into TN, naïve population-closest CD45RA^high^ T_SCM_ (T_CMRA-hi_), CD45RA^low^ T_SCM_ (T_CMRA-low_), CD45RA^negative^ T_SCM_ (T_CMO_), and T_EM_ with CD45RA expression as CD45RA^high^ T_EM_ (T_EMRA-hi_), CD45RA^low^ T_EM_ (T_EMRA-low_) and CD45RA^negative^ T_EM_ (T_EM_) (**Figure S4D**). Longitudinal assessment of T cell naïve/memory composition during expansion showed that, compared to SSC^high^ descendants, SSC^low^ cells-derived progenies generally had a higher portion of T_CMRA-hi_ but less T_CMRA-low_ and other more-differentiated cell subsets (e.g. T_CMO_ and T_EMRA-low_) (**Figures 4D, 4E, and S4E-S4H**). This finding is consistent with a continuously higher expression of co-stimulatory molecules CD27/CD28 that favor cell survival and proliferation (**Figures 4F, S5A, and S5B**) while other phenotypic traits like memory marker CD45RO and inhibition/senescence receptors such as CD57, LAG-3 and TIM-3 were more dynamically variable (**Figures S5A and S5B**).

Functionally, the SSC^low^ cells-derived products responded faster to short period of PKC activation by producing more IL-2 and TNF-α but less IFN-γ (**Figure 4G**). SSC^low^ cells-derived CD4^+^ T cells displayed better compliance with this tendency by expressing the highest amount of early-effector cytokines (e.g. IL-2^+^ and IL-2^+^ TNF-α^+^) and less late-effector cytokines (e.g. IFN-γ^+^, IFN-γ^+^ IL-2^+^ and IFN-γ^+^ TNF-α^+^) (**Figure 4H**). The CD8^+^ T counterpart shows a higher diversity and even conflicts these observations post extensive expansion (e.g. donor 3 at day 11) (**Figure 4I**). Nevertheless, the global cytokine-expressing trait of cell products generated from SSC^low^ group is more naïve-like (**Figures S5C and S5D**) (Gattinoni et al., 2005; Hinrichs et al., 2009), indicating a synergic functionality may happen when CD4^+^ T and CD8^+^ T co-exist in culture system. Superior engraftment and persistence of infused T cells are critical parameters for improved therapeutic index. Extensively expanded T cells from either SSC^high^, SSC^low^ or reconstituted SSC^high+low^ group had similar chemotactic migration to CCL19/CCL21 (**Figure 4J**), in line with their comparable proportion of total T_SCM_ (**Figures 4D and 4E**). However, in response to multiple stimulation by CD3/CD28 engagement, the SSC^low^-derived progenies were endowed with greater capability for multiple expansion under both IL-2 and homeostatic cytokines IL-7/IL-15 (**Figure 4K**), presumably associated with a higher expression of costimulatory markers CD27/CD28 (**Figures 4F, S5A, and S5B**) and underlined epidemic modification favoring cell persistence.

Overall, the SSC^low^ lymphocytes-derived T cell product is discernable from that of SSC^high^ group, the former manifests not only superior proliferation but also earlier phenotypes characteristic of apparently increased T_CMRA-hi_ and enhanced IL-2/TNF-α expression for both CD4^+^ T and CD8^+^ T cells ((**Figures 4L and 4M**).

### Increased T_CMRA-hi_ derived from the SSC^low^ cell cluster resembles central memory precursors

Given the significant accumulation of T_CMRA-hi_ subpopulation that falls in the conventional central memory gate of the cell products derived from SSC^low^ group (**Figures S4D and S1B**), we next sought to define the natural characteristics of this T cell subset by longitudinally determining its phenotypic change during expansion (**Figures 5A–5E**). We showed that T_CMRA-hi_ had a higher expression of canonical memory marker CD45RO than T_N_ but lower than T_CMRA-low_ and T_CMO_ regardless of CD3^+^ CD4^+^ T cells (**Figures 5B and 5C**) or CD3^+^ CD8^+^ T cells (**Figures 5D and 5E**). While all resting central memory T subsets (T_CMRA-hi_, T_CMRA-low_ and T_CMO_ at day 0) presented higher death receptor CD95 (a marker used to distinguish T_SCM_ from T_N_) (Fuertes Marraco et al., 2015; Gattinoni et al., 2011) than the T_N_ subset, they became similar post activation with nearly complete expression of this marker (**Figures 5B–5E**). Notably, the expression of co-stimulatory markers such as CD27 and CD28 was more comparable between T_CMRA-hi_ and T_N_, especially for CD8^+^ T cells (**Figures 5D and 5E**). With regard to inhibitory or senescent factors (e.g. CD57, LAG-3 and TIM-3), CD8^+^ T_CMRA-hi_ and CD8^+^ T_N_ expressed similar but less CD57 than other CD8^+^ T subsets before activation, which however decayed to a dim expression in all subsets post extensive proliferation (**Figure 5D**). In addition, LAG-3 was dramatically increased in activated CD8^+^ T subsets at day 7 and decreased at day 11, during which T_CMRA-hi_ tended to have a higher LAG-3 expression compared to T_N_ but still comparable or even lower than other memory subsets (**Figures 5D and 5E**). These findings suggest that T_CMRA-hi_ is phenotypically distinct from T_N_ and resembles the central memory precursor with naïve-like phenotypic traits (CCR7^+^ CD45RA^high^ CD95^+^ CD28^+^ CD27^+^), which could be taken into consideration in the evaluation of T cell products by different manufacturing protocols.

**Figure 5.**
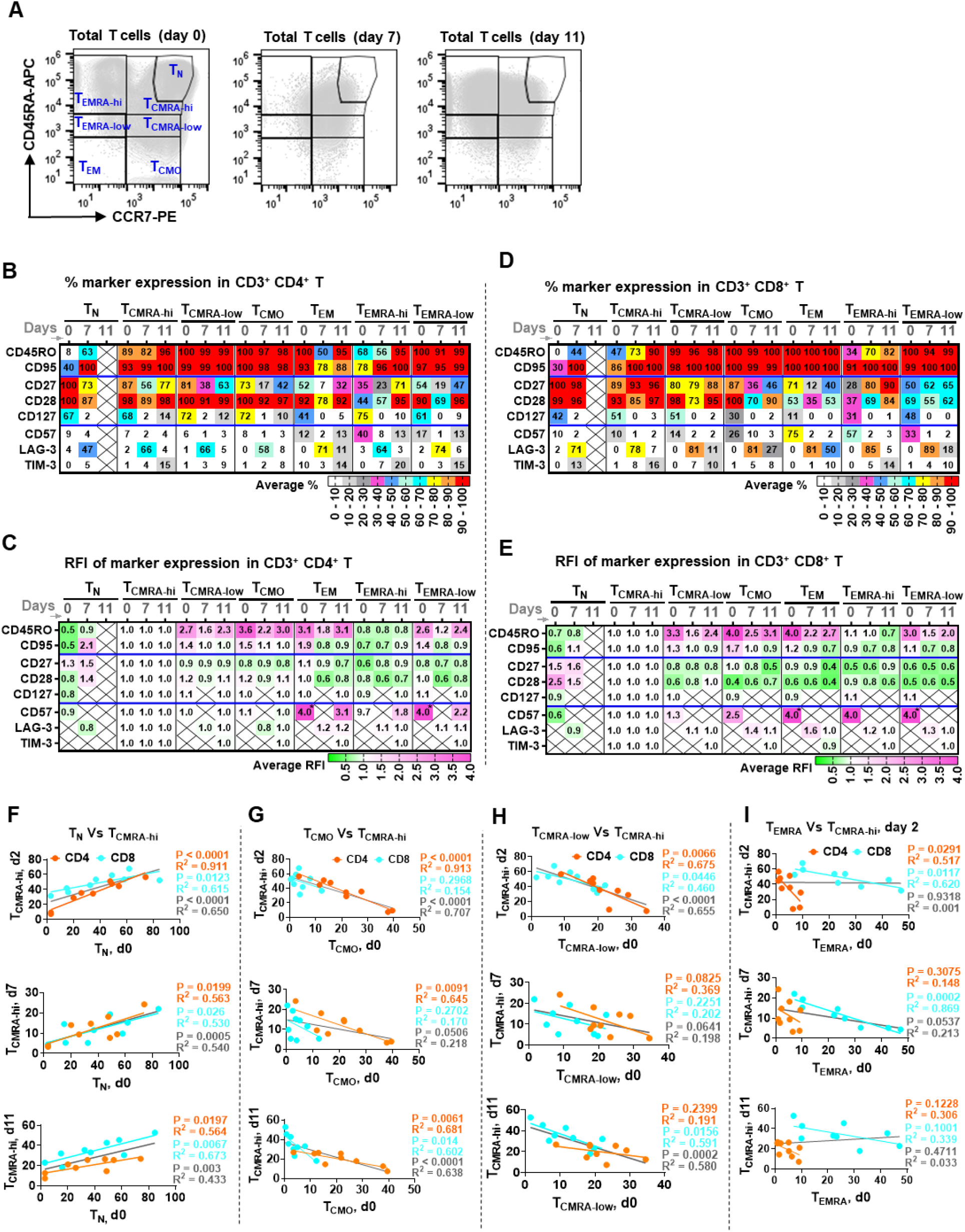
Increased T_CMRA-hi_ derived from SSC^low^ cell cluster resembles central memory precursor. (**A**) Representative plots show the superimposed total T cell populations derived from SSC^hi^, SSC^low^ and SSC^hi^+low cells at day 0, 7, 11. T cell subsets to be analyzed are gated. (**B-E**) Longitudinal assessment of phenotypic change of T_CMRA-hi_ compared to other T cell subsets. Average expression frequency of the indicated markers within CD4^+^ T or CD8^+^ T subsets (**B and E**) and abundance as presented by relative fluorescence intensity (RFI) was reported after normalization to the values of T_CMRA-hi_ for each marker of the same time points respectively (**C and E**). The numbers highlighted by “*” indicate values large than 4. The cross indicates omitted data due to low cell percentages for marker expression identification or insufficient counts for MFI calculation. (**F-I**) Correlation analysis of starting effector/memory composition, including T_N_ (**F**), T_CMO_ (**G**), T_CMRA-low_ (**H**), and total T_EMRA_ (**I**), with the percentage of generated T_CMRA-hi_ among cell products of each time point. Data from the same day from all of the three donors were combined. Each dot represents the percentage of generated T_CMRA-hi_ versus the initial cellular component of its starting cell material (day 0). The solid black line indicates combined linear regression using all the dots in that plot (**F-I**). See also **Figure S4D** for gating method in identifying the dynamic change of T cell lineages during expansion, and **Figures S6A-6C** for individual data of phenotypic marker expression in each T cell subset. **Figures S6D-6G** for correlation analysis of initial cell composition with T_EMRA-hi_ generation.

The burst increase of T_CMRA-hi_ at the beginning of T cell activation (day 2, **Figures 4D and 4E**) is well correlated with the initial proportion of T_N_, and this trend maintains from day 7 to day 11 (**Figure 5F**). In contrast, the existence of more-differentiated CD4^+^ T_CMO_ (**Figure 5G**) and CD4^+^ T_CMRA-low_ (**Figure 5H**) in starting cell materials tend to counteract the generation of CD4^+^ T_CMRA-hi_ despite the effects being varied at different stages post activation. Also, a high number of CD8^+^ T_CMRA-low_ (**Figure 5H**) and CD8^+^ T_EMRA_ (**Figure 5I**) are more likely to reduce the production of CD8^+^ T_CMRA-hi_. However, it is less predictable for CD8^+^ T_EMRA-hi_ generation based on the cell composition before expansion (**Figures S6D-S6G**). Of note, CD4^+^ T_N_ and CD4^+^ T_CMO_ (**Figures S6D and S6E**) do show evidences of contrary correlation with CD4^+^ T_EMRA-hi_ production. There was a sharp decrease of CD8^+^ T_EMRA-hi_ cells post activation in the SSC^high^ group that consists of a larger proportion of differentiated cell subpopulations (**Figures S4E and S4F**). Thus the dynamical accumulation of terminally differentiated T_EMRA-hi_ cells from SSC^low^ groups (**Figures S4F and S4G**) could be partially attributed to a more survival-favoring environment enriched in less-differentiated cell subsets (e.g. T_N_). Additionally, T_EMRA-hi_ displays higher CD27/CD28 expression than the CD45RA-reduced T_EMRA-low_ or T_EM_ subset, which is especially apparent for CD8^+^ T cells during expansion (**Figures 5D and 5E**), suggesting functional potency of this cell subset in cell therapy when alternative co-stimulatory signaling (e.g. CD28 and 4-1-BB) and accessory factors for end-stage effector cells proliferation are involved (Fuertes Marraco et al., 2015; van Leeuwen et al., 2002; Waller et al., 2007). Taken together, these results show that the pre-enrichment of less-differentiated cells simply by side scatter enables the maintenance of naïve-like central memory precursors (T_CMRA-hi_) and accordingly contributes to the slight accumulation of terminally effector-like T_EM_ (T_EMRA-hi_).

### CD4^+^ T cells help is critical for optimum CD8^+^ T cells proliferation

A shaper ratio decrease of CD4/CD8 T cells during expansion was observed for cells from lower side-scatter, which initially have the largest proportion of CD4^+^ T cells and later the highest expansion capacity (**Figure 6A**). This has inspired us to understand the role of CD4^+^ T-CD8^+^ T cells crosstalk in T cell proliferation and functionality shaping despite the absence of specific external antigens (**Figure 6B**).

**Figure 6.**
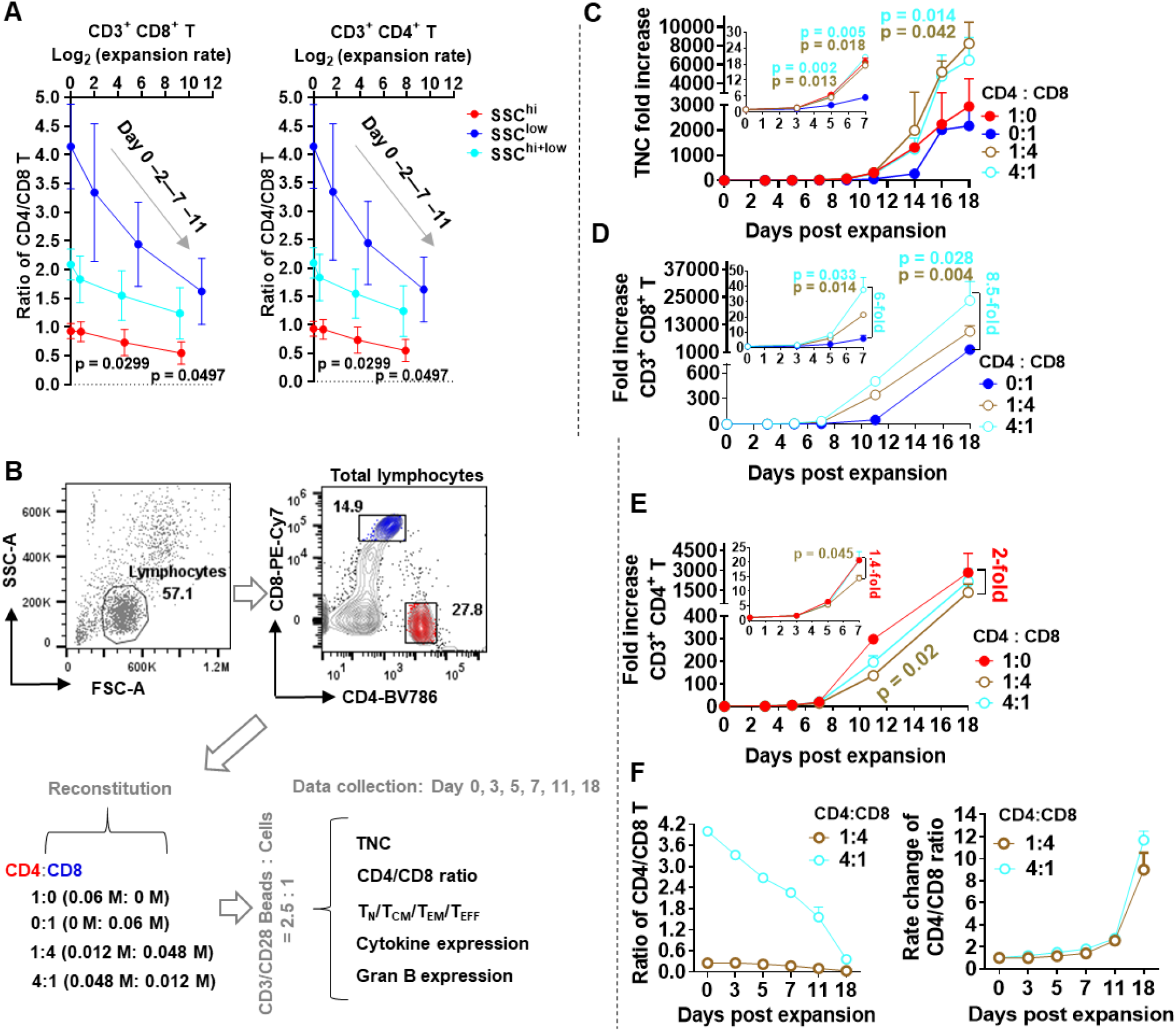
CD4^+^ T help is required for optimal CD8^+^ T expansion. (**A**) Synchronized T cell expansion efficiency with the ratio change of CD4/CD8 T cells starting from cell materials of different side-scattering intensity. Data are representative of three donors as described in Figures 4A-4C. Graphs represent means ± S.E.M (n = 3). Paired two-tailed *t*-test was used for statistical calculation between SSC^hi^ and SSC^low^ group. Error bars indicate mean ± SEM. (**B**) CD4^+^ cells and CD8^+^ cells were sorted and validated to have more than 96% of CD3^+^ CD4^+^ T and CD3^+^ CD8^+^ T cells, respectively. CD4^+^ cells and CD8^+^ cells were reconstituted at the indicated ratio and co-expanded by CD3/CD28 engagement. (**C-E**) Proliferative potential of TNC (**C**), CD3^+^ CD4^+^ T (**D**) or CD3^+^ CD8^+^ T cells (**E**) in condition of CD4-CD8 T cells co-culture. Data of day 5, 7, 11 and 18 represent two independent experiments. Unpaired two-tailed *t-*test compared to culture condition of CD8^+^ T cells only (**C and D**) or CD4^+^ T cells only (**E**). Error bars indicate mean ± SEM. (**F**) Changes in the ratios of CD4^+^ T to CD8^+^ T cells in samples starting with high (4:1) and low (1:4) ratios of CD4 to CD8 T cells. Error bars indicate mean ± SEM.

Co-culture of CD4^+^ T-CD8^+^ T cells favored the global expansion of total T cells (**Figure 6C**). Help from CD4^+^ T cells was required for CD8^+^ T cells maximal proliferation by an average of 6-fold increase at day 7 and 8.5-fold increase at 18 (**Figure 6D**). In turn, CD8^+^ T cells reduced CD4^+^ T cells expansion to a lesser extent by 1.4-fold reduction at day 7 and 2-fold decrease at day 11 and 18 (**Figure 6E**). Thus, the orchestrated growth dynamics within CD4^+^ T and CD8^+^ T cells results in final T cell products with not only different absolute cell numbers (**Figure 6C**) but also discernable CD4^+^ T to CD8^+^ T ratio (**Figure 6F**).

### CD4^+^ T cells promote CD8^+^ T early central memory pool maintenance

By longitudinal assessment of T cell naive/memory composition, we found that optimum maintenance of CD8^+^ T_CMRA-hi_ but less CD8^+^ T_EM_ and CD8^+^ T_EMRA-low_ can be achieved in the presence of CD4^+^ T cells in a ratio-dependent manner (**Figure 7A**). Interestingly, the mild accumulation of CD8^+^ T_EMRA-hi_ was also observed, similar to that of SSC^low^ cells-derived descendants (**Figures S4F and S4G**), indicating a superior culture environment for T cell survival endowed by CD4^+^ T. On the contrary, a large proportion of CD8^+^ T cells tended to counteract CD4^+^ T_CMRA-hi_ generation during the early expansion stages (e.g. day 0-7) (**Figure 7B**). This is reflected by a higher expression of CD28 on CD8^+^ T cells but lower on CD4^+^ T cells throughout the co-culture course (**Figures 7C**, **S7A, and S7B**). Also, the increased number of less-differentiated CD8^+^ T cells generated at day 11 under CD4^+^ T-CD8^+^ T co-expansion is consistent with a greater migration ability to CCL19/CCL21 (**Figure 7D**).

**Figure 7.**
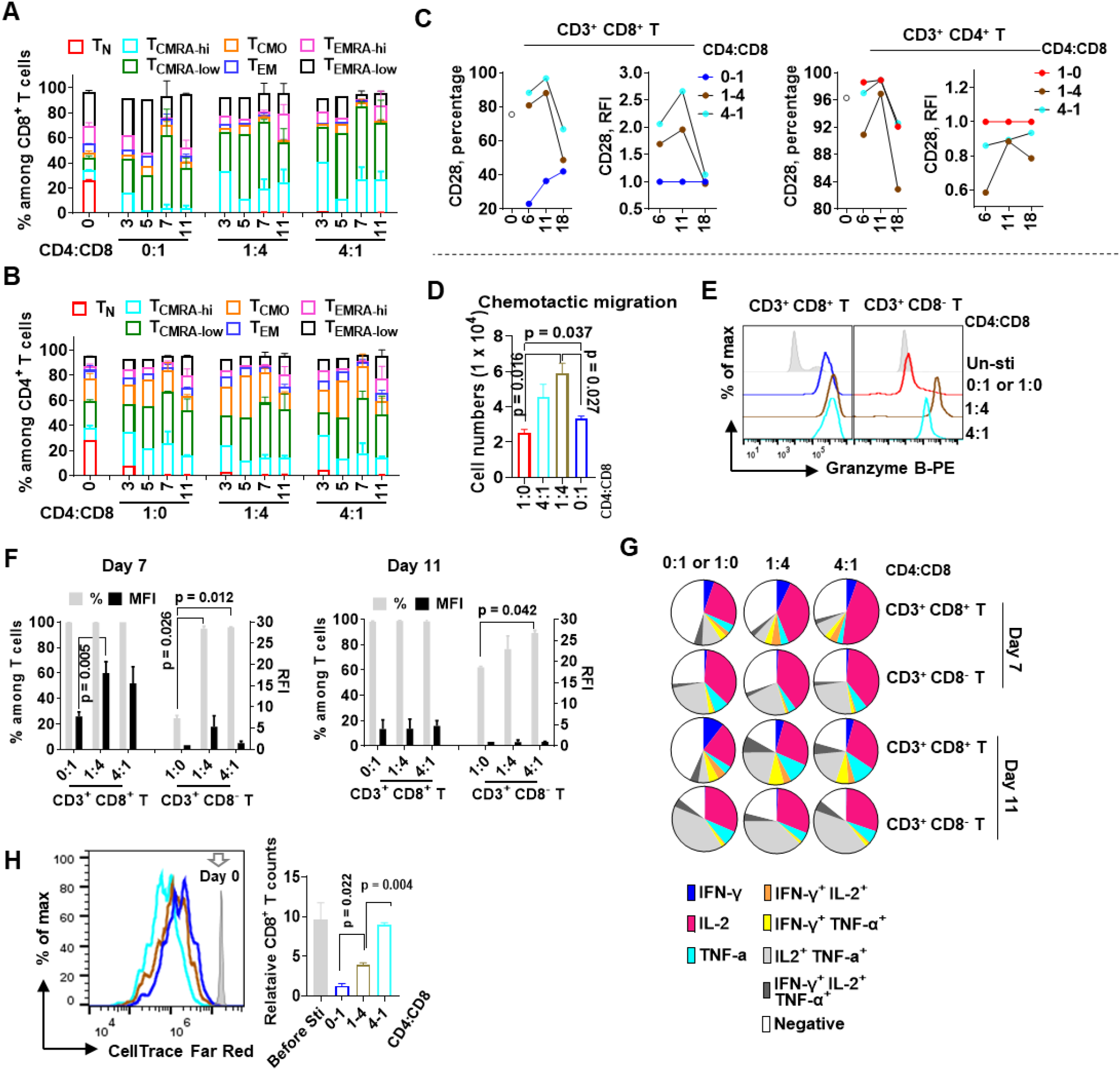
CD4^+^ T cells promote the maintenance of CD8^+^ T early central memory pool. **A and B**) Compositional change of naive/memory subsets in CD3^+^ CD8^+^ T (**A**) or CD3^+^ CD4^+^ T cells (**B**) along the days of expansion. Data for day 0, 7 and 11 are representative of two independent experiments. Error bars indicate mean ± SEM. (**C**) Dynamic expression of CD28 is represented by both percentage and relative fluorescence intensity of T cell products from CD4-CD8 T cells co-expansion. The medium fluorescence intensity (MFI) is normalized to that of CD4^+^ T cells or CD8^+^ T cells cultured separately at the indicated time points. (**D**) CCL21/CCL19-driven migration capacity of T cell generated at day 11 starting from different CD4^+^ to CD8^+^ T cell ratio. Data represents duplicate experiments. Error bars indicate mean ± SEM. (**E and F**) Representative plot of granzyme B-expressing T cells at day 7 post expansion (**E**) and summary of T cells in granzyme B expression from data of two independent experiments (**F**). Error bars indicate mean ± SEM. (**G**) Multifunctional cytokine expression of CD3^+^ CD8^+^ T or CD3^+^ CD4^+^ T cells at day 7 and day 11 post expansion. (**H**) Resting CD4^+^ T cells were sorted and spiked into CellTrace dye-labeled T cell products (day 13) at the indicated ratio and co-stimulated (cells to beads = 1 : 1) for 4 days. Duplicates for each group were performed. Cells were counted at day 4 before collection for anti-CD4-BV786 staining and analysis by flow cytometer. Cell counts were normalized to cell group without newly added CD4^+^ T. Unpaired two-tailed *t*-test between two groups was performed. Error bars indicate mean ± SEM. See also **Figure S7**.

With respect to functional shaping, it is clear that CD4^+^ T and CD8^+^ T cells mutually endowed each other with a higher granzyme B expression during the initial proliferation phase (day 0 - 7, **Figures 7E and 7F**). However, CD8^+^ T cells from all conditional groups became comparable in granzyme B expression at day 11 while a larger portion of CD4^+^ T cells continued to express granzyme B in the presence of CD8^+^ T cells (**Figures 7F**). In addition, CD4^+^ T help contributed to the generation of less-differentiated CD8^+^ T cells characteristic of enhanced IL-2 /TNF-α but reduced IFN-γ production (**Figures 7G**). Instead, CD8^+^ T cells showed only moderate inhibition on the secretion of IL-2/TNF-α by CD4^+^ T cells (**Figures 7G**). Moreover, the inclusion of resting CD4^+^ T cells was able to rescue the expansion potency of extensively differentiated CD8^+^ T cells (**Figures 7H**). Therefore, despite the absence of cognate antigens, help from CD4^+^ T cells is critical for shaping the proliferative capacity and functional potency of CD8^+^ T cells, which should be an important factor to be considered during T cell manufacturing.

## DISCUSSION

TRuST provides a novel method that enables label-free reconstitution of lymphocyte subpopulations using the intrinsic side-scattering property of cells. We demonstrate that less-differentiated T cells are endowed with a lower SSC intensity, which, together with a high ratio of CD4/CD8 T cells, can be effectively enriched and used for optimum T cell production (**Figures 2 and 4**). Furthermore, we address that both naïve T cells and a higher CD4^+^ T to CD8^+^ T proportion are critical for maximal CD8^+^ T cells expansion and optimal CD8^+^ T memory precursors maintenance (**Figures 5–7**).

Naïve T cells have discernable morphological profiles from memory/effector T cells under transmission electron microscope (Dimeloe et al., 2016). Our work demonstrates that this difference can be recaptured simply by using the side-scattering property, which would be more practically meaningful in a high-throughput and cost-effective manner. We showed that SSC is a robust biophysical attribute, the distribution pattern of which presents the intrinsic granularity difference of cellular context and is unaffected by either sample processing or voltage changes applied during data acquisition (**Figures S3A-S3D**). Thus, when applying the gate settings determined using pre-stained cell samples to the unstained cell samples, no gating normalization is required for T cell reconstitution by manual gate selection (**Figures 2 and 4)** and only slight gating correction is required for cell type purification by computational gate search (**Figure 3**). With a single sorting step based on the SSC intensity, lymphocyte populations with high CD4^+^ T to CD8^+^ T ratio are clustered together with younger T cells (e.g. T_N_ and early T_CM_) among the cells of lower SSC. This strategy has been described to meet the demands of current T cell manufacturing procedures in both quantity and quality (**Figures 4, S4, and S5**). A significantly higher yield of T cell with more memory precursors can be achieved from the sorted SSC^low^ cell population. In addition, a comparable amount of T cells with stronger immediate effector function due to the production of late-differentiated effector cells, can also be obtained from the SSC^high^ cluster when compared to unfractionated total lymphocytes. On the other hand, the majority of NK cells can be assigned to SSC^high^ population, which would be useful if antibody-free methods are preferred for NK cells segregation (Hu et al., 2019).

Apheresis materials with high baseline T cell dysfunction (LAG-3^+^/TNF-α^low^ CD8^+^ T) before cell manufacturing was reported to correlate with the non-responding rate of B-CLL patients while a more memory-like functional profile predicts better response (Finney et al., 2019). Similarly, sustained disease remission in B-cell acute lymphoblastic leukemia (B-ALL) patients was associated with memory-like CD27^+^ CD45RO^−^ CD8^+^ T cells before CAR T engineering (Fraietta et al., 2018). Interestingly, we observed that T cell progenies derived from SSC^low^ group have significantly accumulated CD8^+^ T_CMRA-hi_ subpopulation, which is endowed with naïve-like cytokine expression profile (**Figures 4G–4I, S5C, and S5D**) and increased co-stimulatory markers CD27/CD28 but reduced CD45RO/LAG-3 especially during the early expansion phases (**Figures 4F and S5B**). This finding is of great interest for further studies. In addition, tumor histology heterogeneity (e.g. tumor burden and anatomic distribution) and T cell immunosuppression/persistency mechanisms play important roles in cancer immunity and treatment (Majzner and Mackall, 2019). Here we conceive a sequential infusion regimen, that combines highly potential effector cells (SSC^high^-derived) to immediately mitigate tumor load and followed by less-differentiated cells (SSC^low^-derived) to improve long-term tumor surveillance. However, it remains to be validated whether such a treatment regimen would be more efficacious in refractory tumors aimed to maximally alleviate side effects (e.g. cytokine-releasing syndrome and neurotoxicity) while preventing huge tumor burden-caused T cell over-activation/exhaustion. The work has demonstrated its feasibility by producing adequate cell doses but with distinctive cellular quality from one packet of apheresis product.

Great efforts have been put into improving T cell manufacturing processes for better immunotherapy. For example, shortening the time frame of T cell expansion from around two weeks (Jacoby et al., 2019) to a few days (Ghassemi et al., 2018; Lu et al., 2016; Tu et al., 2019) or even without expansion in so called FasT CAR-T technology (Zhang et al., 2019). While more clinical data is required, both strategies have been proposed to maximally maintain the younger T cell subpopulations with enhanced engraftment and persistence. The use of homeostatic cytokine cocktails (e.g. IL-7/IL-15) (Surh and Sprent, 2008) or antigen-specific stimulation to maintain pools of less-differentiated T cells (e.g. T_CM_/T_SCM_) (Cieri et al., 2013; Xu et al., 2014) has also been described. Additionally, CD4^+^ T or CD8^+^ T cells or CD8^+^ T_SCM_ subset can be magnetically pre-enriched and expanded separately in the presence of optimal cytokine cocktails, and co-transfusion of CD4^+^ T and CD8^+^ T cell products has seen preliminary success in both manufacturing and therapeutic studies (Gardner et al., 2017; Shah et al., 2020; Sommermeyer et al., 2016; Turtle et al., 2016). CD4^+^ T help is critical for optimal CD8^+^ T expansion and early central memory pool maintenance in the absence of specific cognate antigens (**Figures 6 and 7**). Hence, it will be of great interest to study how the pre-reconstituted CD4^+^ T/CD8^+^ T cells before expansion may further improve therapeutic efficacy when combined with the defined ratio of CD4/CD8 T cells for infusion. This would be more practically meaningful considering that patients with some tumors or chronic diseases tend to manifest a decreased ratio of CD4/CD8 T cells (Patel et al., 2016; Sommermeyer et al., 2016; You et al., 2009). On the other hand, rapid expansion method (REM), which requires large amount of non-dividing feeder cells such as irradiated allogeneic PBMC or antigen-bearing lymphoblastoid cell line (LCL), has been used to quickly obtain large numbers of therapeutic T cells (Jin et al., 2018; Wang and Riviere, 2015). Our findings would provide an alternative approach to achieve this without the use of additional feeder cells where only a proper ratio of CD4^+^ T to CD8^+^ T cells is adopted for co-expansion. Given these research progresses, our method can be flexibly adapted into those manufacturing workflows.

There are two main advantages for the technical feasibility of TRuST for the preparation of clinical grade T cell products. (1) The lack of expensive staining reagents can significantly reduce cost. By using only one laser (e.g. Blue laser 488 nm) for generating forward-scatter and side-scatter, data-processing burden during real-time sorting can also be simplified. (2) The successful application of good manufacturing practice (GMP)-compliant multiparameter flow cytometry systems in cell therapy (**Table S1**) (Bluestone et al., 2015; Fritsche et al., 2020) and advances in microfluidic systems enabling total lymphocytes enrichment immediately from whole blood in a high-throughput approach by combining acoustic (Ding et al., 2013) and hydrodynamic focusing (Lissandrello et al., 2018; Urbansky et al., 2017). Thus, we envisage the versatility of side scatter-based lymphocytes analysis and reconstitution in these platforms.

### Limitations

Despite the simplicity and ingenuity associated with TRuST, a limitation of this strategy is that it may not be feasible to discriminate differentiated lymphocyte subsets under fast expansion due to confounding effects from enlarged cell size. A possible solution could be the use of longer scattering wavelengths. Also, studies are required in future to test how efficient the SSC-based lymphocyte segregation is when using patient samples which usually manifest abnormal lymphocytes composition and activation statuses. We have demonstrated that it is of adequate sensitivity and pronounced usefulness to segregate the majority of naïve T cells from more-differentiated T cell subsets with controllable CD4/CD8 T cells ratio. We have also showed that, it is feasible to enhance the purification of specific lymphocyte subsets via computational gate design. Nevertheless, it would be more powerful for other applications such as label-free disease diagnosis and highly purified cell selection if the scattering resolution was further improved with interdisciplinary efforts. For example, use of microchannel-based fluidic systems and adoption of adjustable scattering angles (Rossi et al., 2019), inclusion of generic dyes (e.g. MitoTracker) measuring mitochondrial abundance/activity (Baron et al., 2019), or biomarkers possibly surrogating biophysical properties (Tsai et al., 2020; Walsh et al., 2019) to improve lymphocyte subsets identification.

Overall, TRuST is a reliable and effective approach for optimum T cell reconstitution, that will be a significant improvement for current cell manufacturing methods. TRuST is also feasible for the selection of specific lymphocyte subsets for lymphocyte biology studies. This work has provided new insights into the intrinsic light-scattering attributes of different lymphocytes and will pave the way for the improvement and broad application of light property-relied cell identification and separation with interdisciplinary efforts.

## SUPPLEMENTARY INFORMATION

Supplementary information is available for this manuscript.

## ACKNOWLEDGEMENTS

This work was supported by funding from NUS Research Scholarship and Institute for Health Innovation and Technology (iHealthtech), NUS. The authors acknowledge technical support from the Flow Cytometry Laboratory (NUS Medical Sciences Cluster) for cell sorting.

## AUTHOR CONTRIBUTIONS

T.W and L.F.C conceived and designed the study. T.W performed the experiments and data analysis with all other authors’ assistance. T.W and L.F.C wrote the manuscript. Y.H.L, Y.L, and H.J.W reviewed the manuscript and provided feedback. All authors commented on the manuscript and approved the submission.

## METHODS

### Reagents

Apheresis residual blood cones were collected from healthy adult donors from the Health Sciences Authority (HSA), Singapore, with approval from Institutional Review Board and informed consent from donors (NUS-IRB no. H-18-038E). Ficoll^®^ Paque Plus (GE Healthcare, 17-1440-02) was used for peripheral blood mononuclear cells (PBMC) enrichment. Cell culture medium RPMI1640 (Gibco, A1049101) supplemented with heat-inactivated fetal bovine serum (FBS) (Gibco, 10270106) was used for primary cell culture. In some experiments, anti-CD3/CD28 Dynabeads^TM^ (Gibco, 11161D) were used for T cell activation and expansion in the condition of human recombinant IL-2 (Gibco, PHC0021) or where indicated, human recombinant IL-7 plus IL-15. Phorbol ester (PMA) (Sigma-Aldrich, P8139) and ionomycin (Sigma-Aldrich, I9657) were used for cell stimulation to produce cytokines. Cytofix/Cytoperm^TM^ reagent kit (BD Biosciences, 554714) was used for cell fixation and permeabilization before intracellular staining. Flow cytometry antibodies used in this study are listed as follows. Antibodies from BD Biosciences include: CD3-BUV395 (Clone UCHT1; cat #563546), CD4-BV786 (Clone SK3; cat #563877) and CD8-PE-Cy7 (Clone SK1; cat #335787). Antibodies from Biolegend include: CD3-FITC (Clone UCHT1; cat #300406), CCR7-PE (Clone G043H7; cat #353204), CD45RA-APC (Clone HI100; cat #304112), CD45RO-Pacific Blue (Clone UCHL1; cat #304215),CD27-Pacific Blue (Clone M-T271; cat #356413), CD28-FITC (Clone CD28.2; cat #302906), CD95-FITC (Clone DX2; cat #305605), CD127-Brilliant Violet 785 (Clone A019D5; cat #351329), TIM-3-Pacific Blue (Clone F38-2E2; cat #345041), CD57-FITC (Clone HNK-1; cat #359603), LAG-3-Brilliant Violet 785 (Clone 11C3C65; cat #369321) and human TruStain FcX reagent (Cat #422302). Antibodies for intracellular staining include granzyme B-PE (BD Biosciences, GB11; cat #561142), IFN-γ-FITC (Miltenyi Biotec, cat #130-090-433), IL-2-PE (Miltenyi Biotec, cat #130-090-487) and TNF-α-APC (Miltenyi Biotec, cat #130-091-267).

### PBMC isolation and usage

PBMC were isolated from apheresis blood of healthy donors by density gradient centrifugation. To ensure batch-to-batch consistency, the enriched PBMC were counted and cryopreserved at 10 million cells/vial using liquid nitrogen and medium containing 90% FBS and 10% DMSO. For each experiment, the cells were recovered overnight in RPMI1640 supplemented with 10% FBS before downstream experiments.

### T cell differentiation lineages and phenotypes characterization

In this study, two gating methods were used to identify T cell differentiation lineages. For resting T cells, based on the expression level of canonical markers CCR7/CD45RA/CD45RO, they can be broadly classified as naïve (TN, CD45RA^high^ CCR7^high^ CD45RO^−)^, central memory (T_CM_, CD45RA^dim/−^ CCR7^low^), effector memory (TEM, CD45RA^dim/−^ CCR7^−^) and terminally differentiated effector (T_EMRA_, CD45RA^high^ CCR7^−^) (Bohler et al., 2007; Seder and Ahmed, 2003). For T cells undergoing activation and proliferation, an in-depth gating strategy was adapted where the T_CM_ is further dissected into naïve phenotype-closest CD45RA^high^ T_CM_ (T_CMRA-hi_), memory phenotype-closest CD45RA^low^ T_CM_ (T_CMRA-low_) and CD45RA^−^ T_CM_ (T_CMO_), while T_EM_ with CD45RA expression is classified into CD45RA^low^ T_EM_ (T_EMRA-low_) and CD45RA^high^ T_EM_ (T_EMRA-hi_).

To better understand the T cell dynamic changes in their activation/senescent statuses either before or during expansion, the expression profile of a panel of selected cell surface markers was longitudinally assessed, and it includes classical naïve/memory markers CCR7/CD45RA/CD45RO, proliferation and survival-enhancing costimulatory receptors CD27/CD28, and immune activation/inhibition markers CD127/CD95/CD57/LAG-3/TIM-3. To mitigate compensation-induced mutual signaling interferences in multiparameter flow cytometry, the staining panels were designed as follows: panel for dynamic change of naïve/memory composition (CD3-FITC/CD4-BV786/CD8-PE-Cy7/CCR7-PE/CD45RA-APC/CD45RO-V450), panel for co-expression of co-stimulatory markers in naïve/memory subsets (CD3-BUV395/CD8-PE.Cy7/CCR7-PE/CD45RA-APC/CD27-Pacific Blue/CD28-FITC), two panels were designed for immune activation/inhibition markers profiling in naïve/memory T cell subsets (CD3-BUV395/CD8-PE-Cy7/CCR7-PE/CD45RA-APC/CD45RO-Pacific Blue /CD95-FITC/CD127-BV785, or CD3-BUV395/CD8-PE-Cy7/CCR7-PE/CD45RA-APC/TIM-3-Pacific Blue/CD57-FITC/LAG-3-BV785). Pre-blocking step with TruStain FcX (BioLegend) was included for all staining. For some experiments where significant dead cells may be generated such as the first 2-3 days post T cell activation and PMA-based stimulation, LIVE/DEAD viability dye (Invitrogen, L34963) was used to exclude dead cells. CD3-FITC, CD4-BV786, and CD8-PE-Cy7 were from BD Biosciences. All other antibodies used were purchased from BioLegend . Flow cytometry compensation matrices and sample data acquisition were completed in CytoFLEX analyzer (Beckman Coulter) and analyzed by FlowJo software version 10.

### Light-scattering distribution of lymphocytes

To obtain the light-scattering profile of main lymphocyte populations and subpopulations, PBMC were rested overnight, and stained with CD3-BUV395/CD4-BV786/CD8-PE-Cy7 for T cell population identification, or CD3-BUV395/CD56-BV786/CD19-FITC for NK and B cell identification, or CD3-BUV395/CD4-BV786/CD8-PE-Cy7/CCR7-PE/CD45RA-APC for naïve/memory T cells characterization. After washing, the cells were re-suspended in cold PBS before flow cytometry data acquisition. For a label-free approach, the cells were washed twice in cold phosphate buffer (PBS) before suspended in cold PBS and sorted on Moflo Astrios (Beckman Coulter). Sorted cells with different scattering intensity were stained to identify the percentages of CD3^+^ CD4^+^ T/CD3^+^ CD8^+^ T, NK/B cell, or T_N_/T_CM_/T_EM_/T_EMRA_. Where indicated, a panel of selected surface markers was included to probe the detailed activation/senescent phenotypes of sorted cell populations, including CD45RO, CD27, CD28, CD127, CD95, CD57, LAG-3 and TIM-3. The scattering plots were generated by FlowJo software.

### Lymphocytes enrichment by automatic gating search

Computational gating search (GateID) (Baron et al., 2019) was used to find the gate for optional cell type enrichment from a mixed cell population. Briefly, PBMC were recovered overnight and stained with different antibody panels for T cell identification (CD3-BUV395/CD4-BV786/CD8-PE-Cy7), NK/B cells identification (CD3-BUV395/CD56-BV786/CD19-FITC), or naïve/memory T cells characterization (CD3-BUV395/CD4-BV786/CD8-PE-Cy7/CCR7-PE/CD45RA-APC). After washing, flow cytometry data was acquired in CytoFLEX analyzer (Beckman Coulter). The raw readouts corresponding to FSC and SSC parameters for each cell type were exported and input into the GateID package. The yield for each desired cell type (e.g. CD4^+^ T/CD8^+^ T/NK/B/T_N_/T_CM_/T_EM_/T_EMRA_) was set to 5% - 100 % and gating vertices were constantly set to 4. The yield is the count percentage of target cell type within the gating to that of the whole cell population, while the purity indicates the proportion of target cell type to total cell numbers within each gate corresponding to desired yield.

### Cell sorting

For light-scatter based label-free cell sorting from resting PBMC, cryopreserved PBMC were recovered overnight in RPMI1640 supplemented with 10% FBS and washed in cold phosphate buffer (PBS) before suspended in cold PBS and sorted on Moflo Astrios (Beckman Coulter). The excitation wavelength for the generation of side-scatter (488/6-nm filter) and forward-scatter (488/6-nm filter) was 488 nm. Where indicated, a combined excitation by 488 nm and 405 nm was adopted to sort lymphocytes of different side-scatter (488/6-nm and 405/10-nm filter). Side-scatter was measured at an orthogonal angle from the light beam propagation axis while forward-scatter was measured anti-parallel to the incident beam. To make batch-to-batch experiments comparable, the sorting gate was set to equally sort cell populations of different light intensity (SSC^high^/SSC^low^ or FSC^high^/FSC^low^). For antibody staining-based cell sorting, PBMC were pre-blocked with TruStain FcX and stained with CD4-BV786 (BD Biosciences) and CD8-PE-Cy7 (BD Biosciences) for sequential sorting of CD4^+^ cells and CD8^+^ cells on Moflo Astrios. It is important to note that only the cells with the high expression of CD8 were sorted as CD8^+^ T cells because there were some NK cells that also expressed diminished CD8. Also, anti-CD3 was not used for cell sorting experiments where the sorted cells would be subjected to CD3/CD28-engaged stimulation. The sorted cells were validated to achieve enough purity of CD3^+^ CD4^+^ T or CD3^+^ CD8^+^ T cells (> 97%) before reconstituted for co-culture assay.

### Cell stimulation and expansion in vitro

Sorted SSC^low^ and SSC^high^ cells were either expanded alone or reconstituted at a ratio of 1:1 (equal to total lymphocytes within initial cell sample) for co-expansion. The starting cell numbers for each condition were 0.08 million/well in 96-well plate. After 1-hour incubation at 37°C, all cell groups were activated by CD3/CD28 engagement at anti-CD3/CD28 microbeads to cells ratio 3:1 and 5 ng/ml IL-2. Similarly, the sorted CD4^+^ T cells and CD8^+^ T cells were reconstituted at indicated ratios to achieve 0.06 million/well. The ratio of anti-CD3/CD28 microbeads to cells was 3:1 and 5 ng/ml of IL-2 was supplemented. Half of the culture medium was replaced with fresh medium containing 5 ng/ml IL-2 every two days during initial activation/expansion stages (day 3 to day 7) and daily during late expansion phases (day 7 to day 11). During the expansion course, the cell numbers in each group were counted by trypan blue exclusion and aliquoted into approximately 0.1 million/well for all groups to mitigate high cell concentration. Data collection included manual counting of total nucleated cells (TNC), flow cytometry detection of CD4/CD8 T cells, effector/memory composition, and effectors expression (e.g. cytokines and granzyme B) at indicated time points post activation.

Around 5 million expanded T cells at day 11 for each condition were cryopreserved for future usage. In some experiments where cell responsiveness to second stimulation was tested by CD3/CD28 engagement, the pre-expanded T cells at day 11 derived from different SSC groups were thawed and incubated at 37°C overnight. 0.06 million cells were seeded into each well of a 96-well plate and re-stimulated by anti-CD3/CD28 microbeads at beads to cell ratio of 1:1 under 5 ng/ml IL-2 or 5 ng/ml IL-7 and 5 ng/ml IL-15. Total cell numbers were recorded by manual counting and split into approximately 0.1 million/well to minimize high cell concentration. To assess the ability of CD4^+^ T cells to rescue the proliferation capability of extensively pre-expanded CD8^+^ T cells, cell materials derived from SSC^high^ group (day 13) which was enriched in large numbers of terminally differentiated CD8^+^ T cells were labelled with CellTrace Far Red (Invitrogen, C34564) following the manufacturer protocol. The labelled T cells and sorted resting CD4^+^ T cells of the same donor were reconstituted at indicated ratios and activated by anti-CD3/CD28 microbeads supplemented with 5 ng/ml IL-2. After 4 days, total cell numbers were counted and stained with CD4-BV786 before analysis on flow cytometer to determine CD8^+^ T cells proliferation.

### Functionality in effecting factors expression

To determine the immune response of T cells in cytokine and granzyme B expression, residual anti-CD3/CD28 microbeads were removed from expanded T cells at day 7 or day 11 before being re-stimulated in the presence or absence of 50 ng/ml PMA and 1 μg/ml ionomycin for an additional 2.5 hours. Cells were collected and labelled with LIVE/DEAD viability dye before further blocking with TruStain FcX. Surface markers CD3-BUV395 and CD8-PE-Cy7 were used to characterize T cell populations. After fixation/permeabilization treatment, each sample was divided into two tubes for either cytokine detection with IFN-γ-FITC (Miltenyi Biotec), IL-2-PE (Miltenyi Biotec) and TNF-α-APC (Miltenyi Biotec) or granzyme B measurement with granzyme B-PE (BD Biosciences) where isotype control antibodies were properly included. Flow cytometry data were acquired in CytoFLEX analyzer (Beckman Coulter) and analyzed by FlowJo software. Notably, the CD4 marker has a substantial downregulation post PMA stimulation, thus the CD3^+^ CD8^−^ T cells were gated as CD3^+^CD4^+^ T cells instead for data presentation.

### Migration assay

*In vitro* chemotactic migration was used to evaluate the cells lymphnode-homing capacity. Briefly, CCL19 (300 ng/ml) and CCL21 (300 ng/ml) (PeproTech) in 100 μl culture medium were added to the lower chamber of 96-well transwell plate (5 μm porosity) (Corning, CLS3388). For control group, only culture medium without chemokines was added. 0.1 million expanded T cells (day 11) derived from SSC^high^, SSC^low^ and reconstituted SSC^high+low^ groups, or from experiments using CD4^+^ T-CD8^+^ T co-culture were seeded into the upper chamber in 80 μl culture medium. Each condition was conducted in duplicates. After five hours, the numbers of cells migrating to the lower chamber were counted by trypan blue exclusion. For migration capacity, the counts subtracted from the migrated cell numbers in control group were presented, correspondingly.

### Principal component analysis

For each cell sample derived from sorted SSC^high^, SSC^low^ or reconstituted SSC^high+low^ population, the cell composition (T_N_, T_CMRA-hi_, T_CMRA-low_, T_CMRA-low_, T_CMO_, T_EM_, T_EMRA-hi_, and T_EMRA-low_) and multifunctional cytokine expression profile (IFN-γ^+^ only, IL-2^+^ only, TNF-α^+^ only, IFN-γ^+^ IL-2^+^, IFN-γ^+^ TNF-α^+^, IL-2^+^ TNF-α^+^, and IFN-γ^+^ IL-2^+^ TNF-α^+^) were integrated together for PCA calculation using the program embedded in Seurat package (Stuart et al., 2019). For PCA plotting, the same cell subsets from all groups with percentage less than 4% were omitted. The data for day 7 and day 11 were calculated separately. For data at day 7, the IFN-γ^+^ IL-2^+^/IFN-γ^+^ TNF-α^+^/IFN-γ^+^ IL-2^+^ TNF-α^+^ were combined as “IFN^+^ Multi” due to relatively lower percentage of these cells in some donors expressing double positive cytokines.

### Statistical methods

All values with error bars are reported as mean ± SEM. Statistical analysis was performed using GraphPad Prism. Unpaired two-tailed *t*-test was used to evaluate statistical significances between two groups. Where indicated, paired two-tailed *t*-test was used to compare the differences between two groups that combined data of different donors together.

### DATA AND CODE AVAILABILITY

The data that support the findings of this study can be made available by the corresponding author. There is no customized code for this work. The set of adjustable parameters in GateID program has been indicated in the Methods section.

