## Supplemental Fig. 1-7 & Table 1 for "Label-free lymphocytes reconstitution using side scatter for optimal T cell manufacturing"

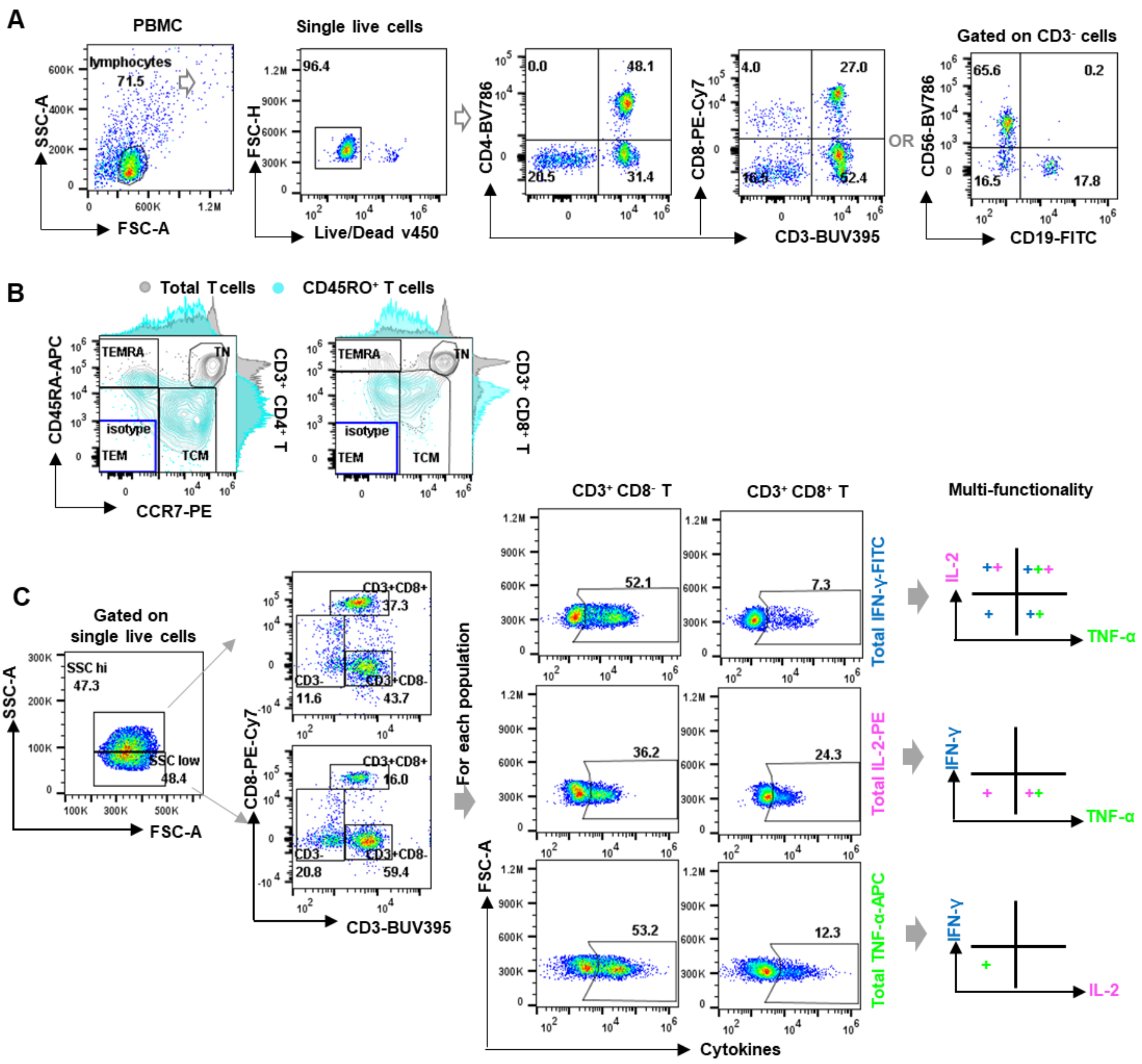


**Figure S1. Gating methods to identify lymphocyte populations and cytokine-expressing potency, Related to Figure 2**

(**A**) Gating strategies to identify CD3$^+$ CD4$^+$ T, CD3$^+$ CD8$^+$ T, NK and B cells.

(**B**) Characterization of T cell naïve/memory lineages including $T_N$, $T_{CM}$, $T_{EM}$, and $T_{EMRA}$ based on the expression of canonical markers CCR7/CD45RA/CD45RO. Blue line represents CCR7/CD45RA isotype gate.

(**C**) Representative gating method for multifunctional cytokine expression analysis of lymphocytes with different SSC intensity. For unstained cell sample, the cells were pre-sorted based on the SSC intensity into $SSC^{high}$ and $SSC^{low}$ clusters before subsequent activation and cytokine detection. The percentage of multifunctional cytokine expressing cells among $CD3^+$ $CD8^-$ T or $CD3^+$ $CD8^+$ T cells was further normalized based on the total expression of each cytokine.

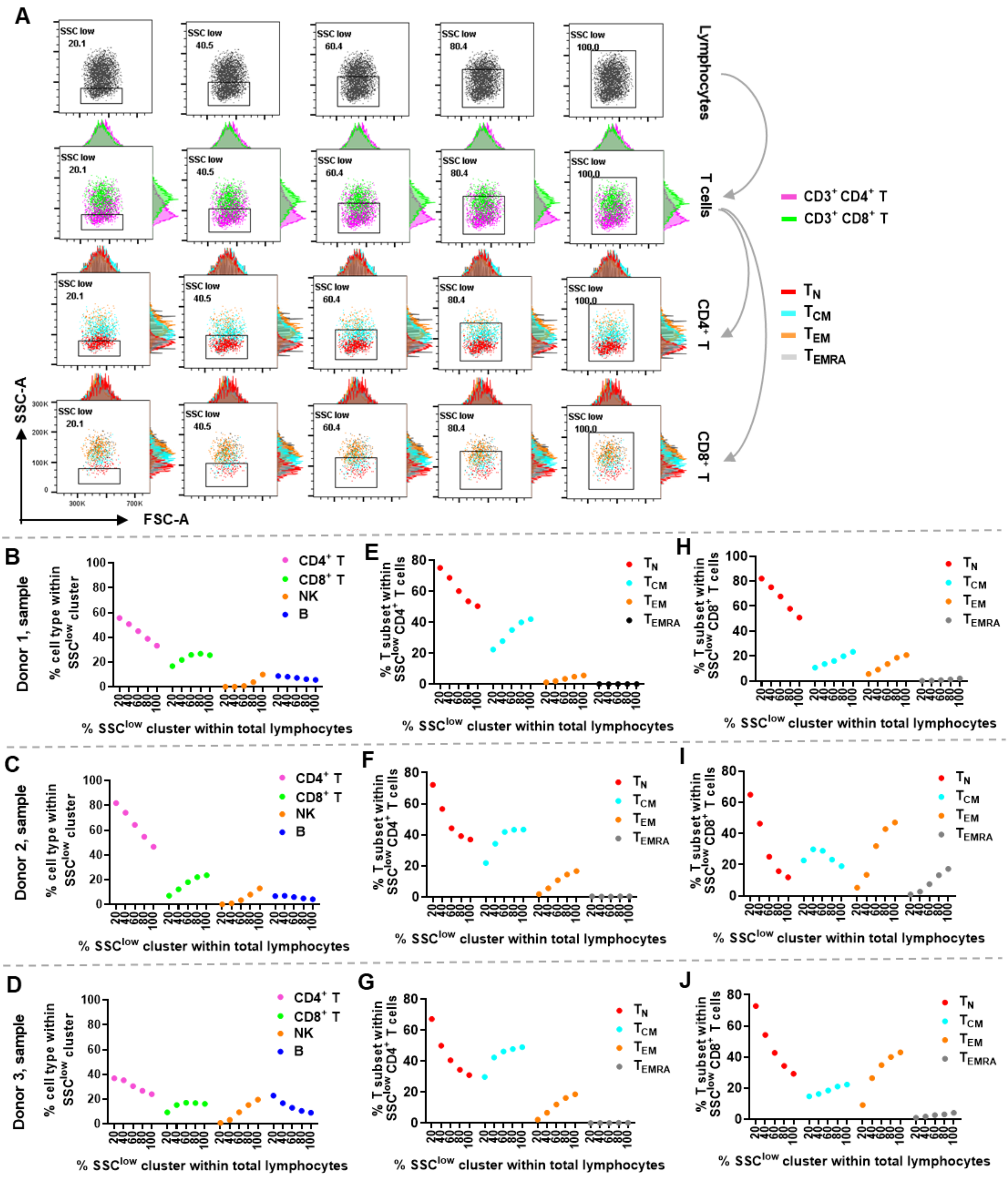

**Figure S2. Manual gate selection for T cell reconstitution, Related to Figure 2**

(**A**) A serial of gates based on the SSC intensity were set to recover various amount of SSC$^{low}$ lymphocytes, representing 20% to 100% of the total lymphocytes analyzed. The scattering distribution of CD3⁺ CD4⁺ T, CD3⁺ CD8⁺ T, and T cell naïve/memory lineages within the gated

$SSC^{low}$ cluster was displayed.

(**B-J**) Shown are the percentage of $CD3^+$ $CD4^+$ T, $CD3^+$ $CD8^+$ T, NK and B cells (**B-D**), or T cell naïve/memory subsets in $CD3^+$ $CD4^+$ T (**E-G**) and $CD3^+$ $CD8^+$ T cells (**H-J**) among the gated $SSC^{low}$ cluster (n = 3).

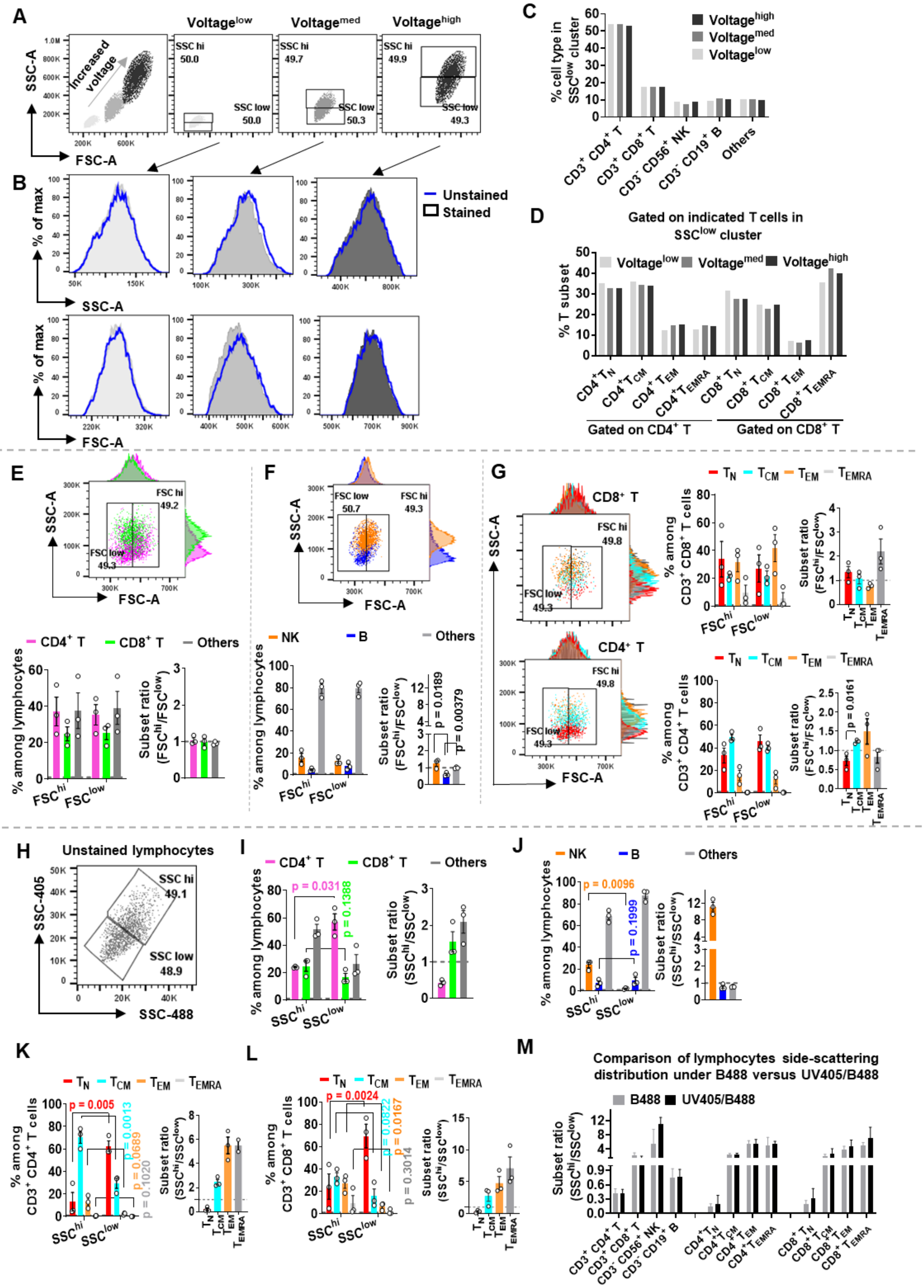

A
Voltage^low
Voltage^med
Voltage^high
SSC-A
FSC-A
Increased voltage
SSC hi 50.0
SSC low 50.0
SSC hi 49.7
SSC low 50.3
SSC hi 49.9
SSC low 49.3
B
Unstained
Stained
% of max
SSC-A
FSC-A
C
% cell type in SSC^low cluster
CD3+ CD4+ T
CD3+ CD8+ T
CD3- CD56+ NK
CD3- CD19+ B
Others
D
Gated on indicated T cells in SSC^low cluster
% T subset
Gated on CD4+ T
Gated on CD8+ T
E
F
G
FSC hi
FSC low
CD4+ T
CD8+ T
Others
NK
B
% among lymphocytes
Subset ratio (FSC^hi/FSC^low)
p = 0.0189
p = 0.00379
p = 0.0161
% among CD3+ CD8+ T cells
% among CD3+ CD4+ T cells
H
Unstained lymphocytes
SSC-405
SSC-488
SSC hi 49.1
SSC low 48.9
I
J
Subset ratio (SSC^hi/SSC^low)
p = 0.031
p = 0.1388
p = 0.0096
p = 0.1999
K
L
p = 0.005
p = 0.0013
p = 0.0689
p = 0.1020
p = 0.0024
p = 0.0822
p = 0.0167
p = 0.3014
M
Comparison of lymphocytes side-scattering distribution under B488 versus UV405/B488
B488
UV405/B488

**Figure S3. Side scatter is a robust parameter for lymphocytes segregation, Related to Figure 2**

(**A-D**) PBMC was pre-stained with fluorescence antibodies and run on flow cytometer under different voltages for both FSC and SSC (**A**). Control cells instead were unstained but subjected to the same sample processing steps. Shown is the scattering profile of total lymphocytes with or without antibody staining (**B**). The percentage of $CD3^+$ $CD4^+$ T, $CD3^+$ $CD8^+$ T, NK and B cells in gated $SSC^{low}$ cluster (**C**), as well as the naive/memory T cell subsets in $CD4^+$ T and $CD8^+$ T lymphocyte among the gated $SSC^{low}$ cell population (**D**) were compared under different voltages as applied for data acquiring.

(**E-G**) Alternative gating method based on the FSC intensity was used to divide total lymphocytes into $FSC^{hi}$ and $FSC^{low}$ clusters as illustrated in the dot plots**.** The percentage of $CD3^+$ $CD4^+$ T/$CD3^+$ $CD8^+$ T cells (**E**), NK/B cells (**F**), and $T_N$/$T_{CM}$/$T_{EM}$/$T_{EMRA}$ cell subsets (**G**) within the $FSC^{hi}$ or $FSC^{low}$ cluster were summarized (histograms) (n = 3). Data are presented as means ± S.E.M (n = 3). Paired two-tailed *t*-test between $SSC^{low}$ and $SSC^{hi}$ group was performed. Error bars indicate mean ± SEM.

(**H-M**) Use of double side-scattering gate (**H**) for segregation of unstained lymphocyte populations. Shown are $CD4^+$ T and $CD8^+$ T lymphocyte composition (**I**), B and NK proportion (**J**) in sorted $SSC^{hi}$ and $SSC^{low}$ cell populations (n = 3), or naive/memory T cell subsets in $CD4^+$ T (**K**) and $CD8^+$ T lymphocyte (**L**) within the sorted $SSC^{hi}$ and $SSC^{low}$ cell populations, respectively (n = 3). (**M**) The data from **Figures S3I-S3L** and **Figures 2D-2G** were compiled together to show the efficiency of lymphocytes reconstitution sorted under either blue laser (B488) or combined with UV laser (UV405). A two-tailed, paired *t*-test between two groups was performed (**E-I**). Error bars indicate mean ± SEM.

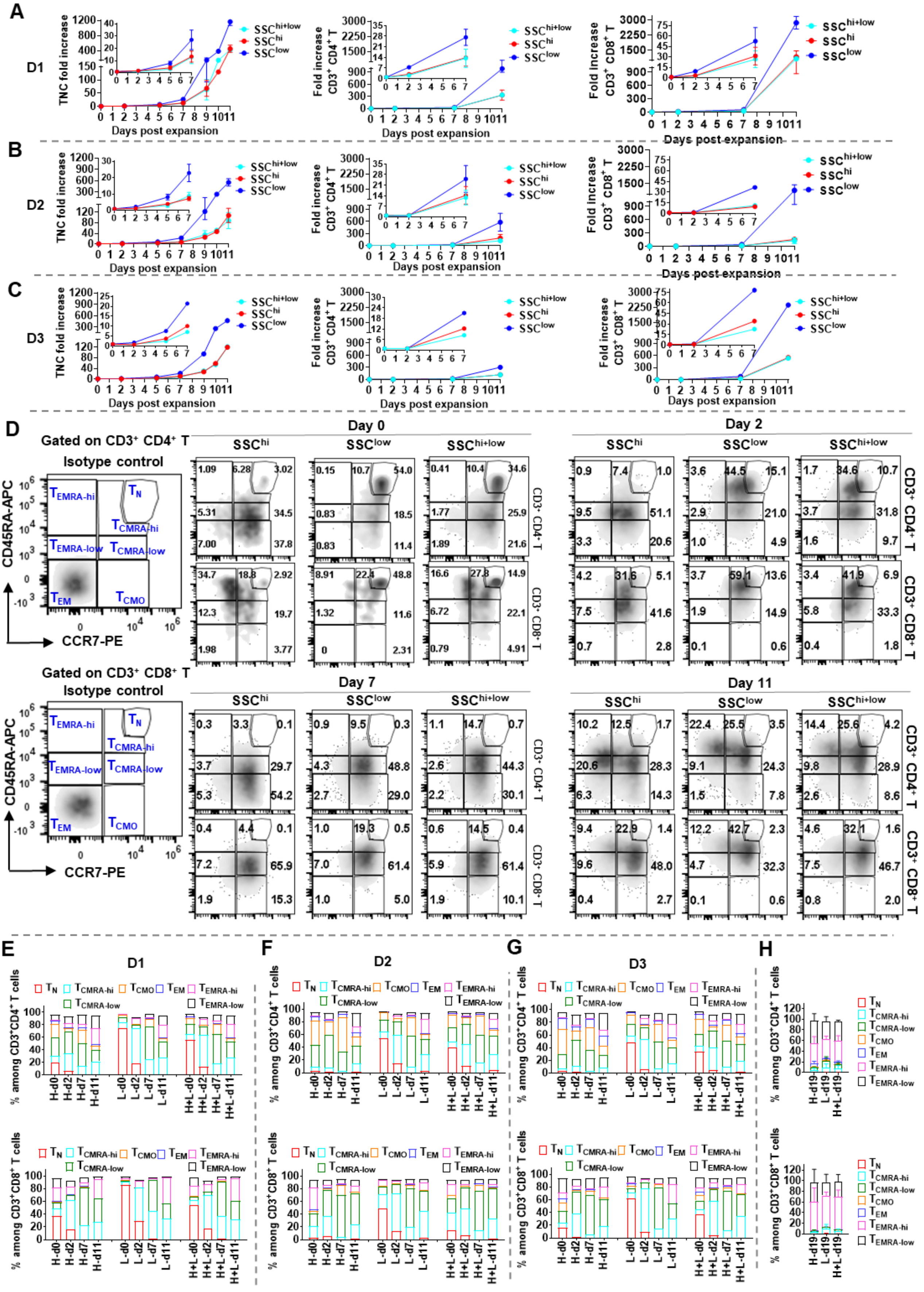

**Figure S4. Dynamic expansion and phenotype shaping of side-scatter reconstituted lymphocytes from individual donors, Related to Figure 4**

(**A-C**) Expansion efficiency of lymphocytes derived from $SSC^{hi}$, $SSC^{low}$ , or reconstituted $SSC^{hi+low}$ cells at a ratio of 1:1 as represented by TNC fold increase (**A**), $CD3^+$ $CD4^+$ T (**B**) and $CD3^+$ $CD8^+$ T (**C**) fold increase. Two independent experiments for donor 1 and donor 2 and one experiment for donor 3 are shown. Error bars indicate mean ± SEM.

(**D**) Gating strategy for in-depth analysis of compositional change of naïve/memory T cells during expansion.

(**E-G**) Composition change of naive/memory T subsets in $CD3^+$ $CD4^+$ T or $CD3^+$ $CD8^+$ T cells for each donor along the expansion procedure. Two independent experiments for donor 1 (**E**) and donor 2 (**F**) at day 0, 7 and 11 and one experiment for donor 3 (**G**) are shown.

(**H**) T cell naïve/memory composition at day 19 post expansion (n = 2).

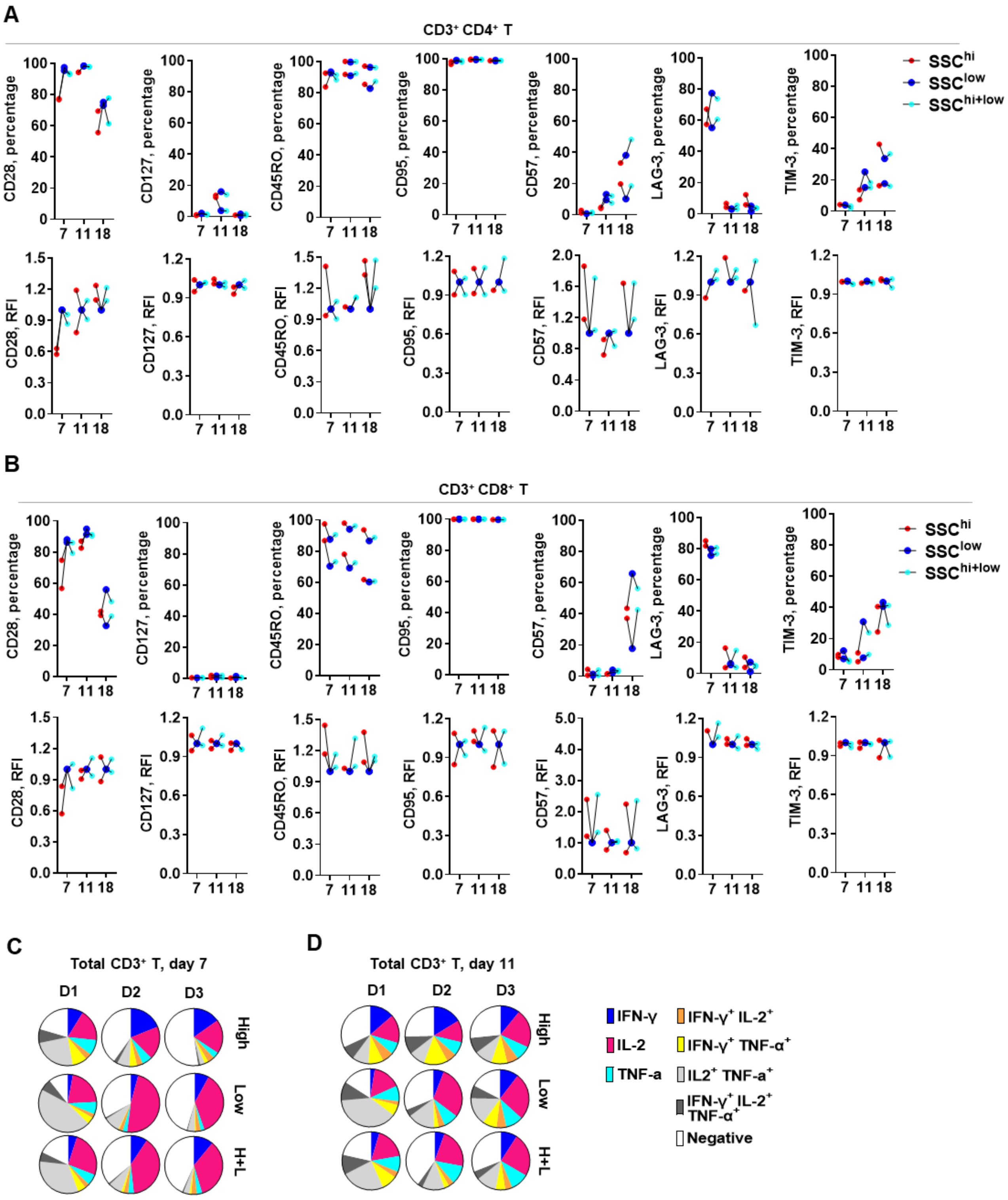


**Figure S5. Phenotype and functionality of of reconstituted lymphocytes post activation, Related to Figure 4**

(**A and B**) Dynamic expression of a panel of phenotypic markers in total CD3⁺ CD4⁺ T (**A**) and

$CD3^+$ $CD8^+$ T cells (**B**) at day 7, 11 and 18 post expansion (n = 2).

(**C and D**) Multifunctional cytokine expression of total $CD3^+$ T cells at day 7 (**C**) and day 11 (**D**) post expansion (n = 3) with 2 independent experiments for donor 1 and 2.

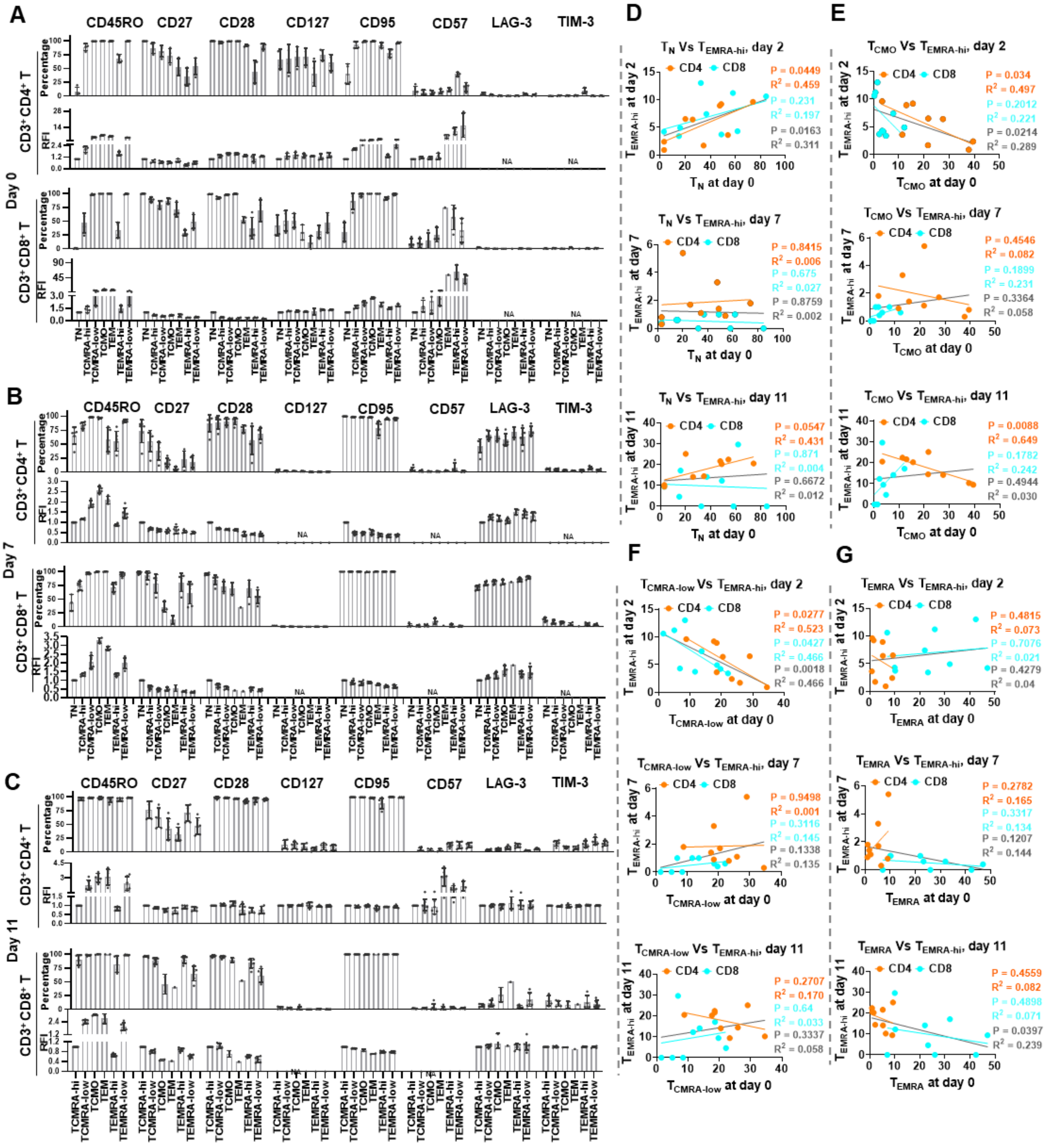

**Figure S6. CD4⁺ $T_{EMRA\text{-}hi}$ but not CD8⁺ $T_{EMRA\text{-}hi}$ in T cell products is predictable from initial proportion of $T_N$ and $T_{CMO}$, Related to Figure 5**

(**A-C**) Phenotypic characterization of $T_{CMRA\text{-}hi}$ subset compared to $T_N$, $T_{CMRA\text{-}low}$, $T_{CMO}$, $T_{EM}$, $T_{EMRA\text{-}low}$ and $T_{EMRA\text{-}hi}$. Flow cytometry data of expanded T cells at day 0 (**A**), day 7 (**B**) and day 11 (**C**).

The relative abundance is represented as relative fluorescence intensity (RFI) after normalization to that of $T_N$ for day 0 and day 7, respectively (**A and B**) or to that of $T_{CMRA-hi}$ for day 11 due to nearly undetectable $T_N$ (**C**). For each T cell subset, a total of 4 readouts (day 0) and 6 readouts (day 7 or day 11) from $SSC^{hi}$, $SSC^{low}$ and $SSC^{hi+low}$ groups (n = 2) were combined and reported after removing cell counts collected less than 50. “NA” indicates omitted data due to generally low cell counts for MFI calculation. Error bars indicate mean ± SEM.

(**D-G**) Correlation analysis of starting effector/memory composition including $T_N$ (**D**), $T_{CMO}$ (**E**), $T_{CMRA-low}$ (**F**, or $T_{EMRA}$ (**G**) with the percentage of generated $T_{EMRA-hi}$ among cell products corresponding to each time point along the days of expansion. Data of the same day from three donors were combined. Each dot represents the percentage of generated cell subset corresponding to the indicated cell component of its starting cell material. The solid black line indicates combined linear regression using all the dots in that plot. Note that the total $T_{EMRA}$ at day 0 is taken for analysis due to the extremely low percentage of $T_{EMRA-hi}$ among $CD4^+$ T cells.

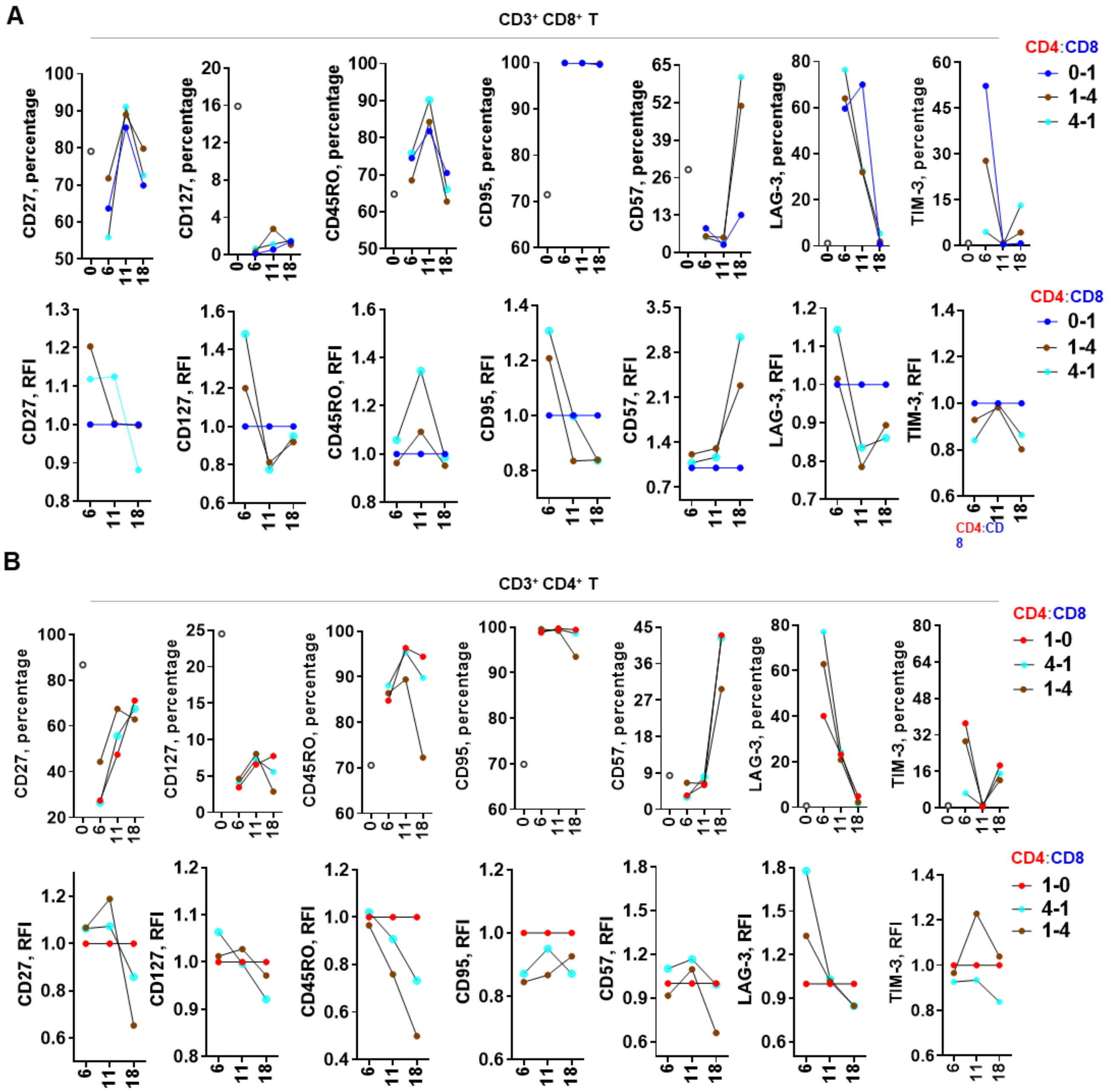


**Figure S7. Expression level of selected phenotypic markers of reconstituted lymphocytes during expansion, Related to Figure 7**

(**A and B**) Dynamic expression of a panel of phenotypic markers in CD3$^{+}$ CD4$^{+}$ T (**A**) or CD3$^{+}$ CD8$^{+}$ T cells (**B**) at day 6, 11 and 18 post CD4-CD8 T co-expansion. The relative fluorescence intensity (RFI) is calculated by normalizing the medium fluorescence intensity (MFI) of each marker under CD4-CD8 co-culture conditions to that of CD4$^{+}$ T cells or CD8$^{+}$ T cells culture at the indicated time points.

**Table S1. Technical feasibility analysis of side-scattering relied cell sorting for therapeutic purpose, Related to Figure 1**

| GMP-compliant cell sorter | Sorting system | Multi-parameters | Sorting channel width | Other features | Flow rate & sort rate | Loading cell number | Estimated time for $SSC^{low}$ lymphocytes enrichment[a] | Estimated T cell numbers post expansion | Cells per dose tested in clinic (body weight: 80 kg) |
|---|---|---|---|---|---|---|---|---|---|
| MACS Quant® Tyto® cell sorter | Disposable, closed cartridge integrated with microchip; 2-way sorting | 3 lasers (405 nm, 488 nm, 638 nm); Back scatter; Side scatter | 25×50 µm | High-speed; No droplets; No charge; Low pressure (< 3 psi); | Up to 55,000 cells/s; 30,000 valve actuations/s | Up to $5×10^7$ cells/ml in 10 ml buffer per cartridge | $6×10^7$ targets in 3 hours by two sort processes starting from $5×10^8$ PBMC; Or 4.8 $×10^6$ targets in 2.5 hours by single running starting from $2×10^7$ PBMC[b] | Day 0: $6×10^7$ $SSC^{low}$ lymphocytes<br><br>Day 7: $1.2×10^9$ (~ 20-fold increase);<br><br>Day 11: $3.6×10^{12}$ (~ 600-fold increase) | Without expansion (FasT CAR-T) (Zhang et al., 2019): ~ $4×10^6$ - $1.2×10^7$<br><br>7-21 days expansion (Jacoby et al., 2019) : ~ $4×10^7$ - $2.4×10^8$<br><br>Solid tumor (Zhang et al., 2017): ~ $10^7$ - $10^9$ |
| Sony FX500 cell sorter | Jet-in-air, replaceable fluidics system; 2-way sorting | 3 lasers (488 nm, 561 nm, 638 nm); Back scatter; Forward scatter | 70-130 µm nozzle integrated with microchip | High-speed; Droplet; Charged; Tens of psi; | Up to 30,000 cells/s; | Unlimited (sample in tube) | $6×10^7$ targets in 3 hours by single running (20,000 events/s) starting from $2×10^8$ PBMC[c] | | |

[a]For PBMCs post density gradient centrifugation, the total lymphocyte population approximately accounts for 70-90%. For PBMCs with high proportion of monocytes contamination, plate adherence or counter-flow elutriation can be used to improve lymphocytes composition.

[b]The estimated yields of $SSC^{low}$ lymphocytes is calculated based on scientific reports from Miltenyi Biotec website. For the processing of $5×10^8$ PBMC, it took around 3 hours to complete sample running by a two sort processes: initial bulk sorting ($5×10^8$ cells/ml) for 135 min + second purify sorting ($2×10^6$ cells/ml) for additional 45 min. By this, a final yield of target cells is around 30% with > 90% purity. For the processing of smaller cell materials (e.g. $2×10^7$ PBMC), it took around 2.5 hours to complete sample running by a single running ($2×10^6$ cells/ml) to achieve average yield of around 60% (some up to 80%) with > 90% purity.

[c]The estimated yields of $SSC^{low}$ lymphocytes is calculated based on reporting from Sony Biotechnology website and a comparison between MACS Quant Tyto with Sony FX500 (Radtke et al., 2019). For cell sample at $2×10^7$ cells/ml, using the 70 µm sorting chip, a flow rate of 20,000 events/s can be used to sort target cells with a purity of > 99% and efficiency of > 75%.
